# Cystathionine-β-synthase is essential for AKT-induced senescence and suppresses the development of gastric cancers with PI3K/AKT activation

**DOI:** 10.1101/2021.07.04.451041

**Authors:** Haoran Zhu, Keefe T. Chan, Xinran Huang, Shaun Blake, Anna S. Trigos, Dovile Anderson, Darren Creek, David P. De Souza, Xi Wang, Caiyun Fu, Metta Jana, Elaine Sanij, Richard B Pearson, Jian Kang

## Abstract

Hyperactivation of oncogenic pathways downstream of RAS and PI3K/AKT in normal cells induces a senescence-like phenotype that acts as a tumor-suppressive mechanism that must be overcome during transformation. We previously demonstrated that AKT-induced senescence (AIS) is associated with profound transcriptional and metabolic changes. Here, we demonstrate that human fibroblasts undergoing AIS display increased Cystathionine-β-synthase (CBS) expression and consequent activation of the transsulfuration pathway controlling hydrogen sulfide (H2S) and glutathione (GSH) metabolism. Activated transsulfuration pathway during AIS maintenance enhances the antioxidant capacity, protecting senescent cells from ROS-induced cell death via GSH and H2S. Importantly, CBS depletion allows cells that have undergone AIS to escape senescence and re-enter the cell cycle, indicating the importance of CBS activity in maintaining AIS. Mechanistically, we show this restoration of proliferation is mediated through suppressing mitochondrial respiration and reactive oxygen species (ROS) production and increasing GSH metabolism. These findings implicate a potential tumor-suppressive role for CBS in cells with inappropriately activated PI3K/AKT signaling. Consistent with this concept, in human gastric cancer cells with activated PI3K/AKT signaling, we demonstrate that CBS expression is suppressed due to promoter hypermethylation. CBS loss cooperates with activated PI3K/AKT signaling in promoting anchorage-independent growth of gastric epithelial cells, while CBS restoration suppresses the growth of gastric tumors *in vivo*. Taken together, we find that CBS is a novel regulator of AIS and a potential tumor suppressor in PI3K/AKT-driven gastric cancers, providing a new exploitable metabolic vulnerability in these cancers.

## Introduction

Hyperactivation of oncogenic pathways such as RAS/ERK or PI3K/AKT can cause cellular senescence in non-transformed cells, termed oncogene-induced senescence (Serrano et al., 1997; Zhu et al., 2020). In addition to the well-studied RAS-induced senescence (RIS), we and others have demonstrated that hyperactivation of PI3K/AKT signaling pathway causes a senescence-like phenotype, referred to as AKT-induced senescence (AIS) or PTEN loss-induced cellular senescence (PICS) (Alimonti et al., 2010; Astle et al., 2012; Chan et al., 2020; Jung et al., 2019). AIS is characterized by common senescence hallmarks including cell cycle arrest, a senescence-associated secretory phenotype (SASP), global transcriptional changes and metabolic hyperactivity (Chan et al., 2020). Distinct from RIS, AIS does not display either p16 upregulation, a DNA damage response or senescence-associated heterochromatin foci. Instead, AIS is associated with elevated p53 expression through increased mTORC1-dependent translation and reduced human double minute 2 (HDM2)-dependent destabilization (Astle et al., 2012). Disruption of the critical mechanisms regulating maintenance of oncogene-induced senescence can lead to tumorigenesis (Braig et al., 2005; Chen et al., 2005; Collado et al., 2005). Therefore, understanding the molecular mechanisms that regulate AIS and how they are subverted will provide opportunities to identify therapeutic strategies for suppressing PI3K/AKT-driven cancer development.

We identified 98 key regulators in a whole-genome siRNA AIS escape screen and validated a subset of these genes in the functional studies to confirm their role in AIS maintenance (Chan et al., 2020). Intriguingly, 11 genes were associated with the regulation of metabolism, suggesting that an altered metabolism could be integral for maintaining AIS. In particular, the cystathionine-β-synthase (CBS) was ranked as one of the top metabolic gene candidates with loss of expression leading to AIS escape (Chan et al., 2020), but how it does so is not known.

CBS is an enzyme involved in the transsulfuration metabolic pathway. CBS converts homocysteine (Hcy), a key metabolite in the transmethylation pathway, to cystathionine which is subsequently hydrolysed by cystathionine gamma-lyase (CTH) to form cysteine, the crucial precursor for glutathione (GSH) production (Fig. 1). CBS also catalyses the production of hydrogen sulfide (H2S), a diffusible gaseous transmitter that modulates mitochondrial function and cellular bioenergetics (Szabo et al., 2013; Szabo et al., 2014); exerts antioxidant effects through inhibition of reactive oxygen species (ROS) generation and lipid peroxidation (Wen et al., 2013); and stimulates antioxidant production via sulfhydration of key proteins involved in antioxidant defence such as Keap1 and p66Shc (Paul & Snyder, 2012; Yang et al., 2013). Thus, CBS acts through control of Hcy, H2S and GSH metabolism and exerts diverse biological functions including regulating DNA methylation, mitochondrial respiration and redox homeostasis (Zhu et al., 2018).

**Figure 1.**
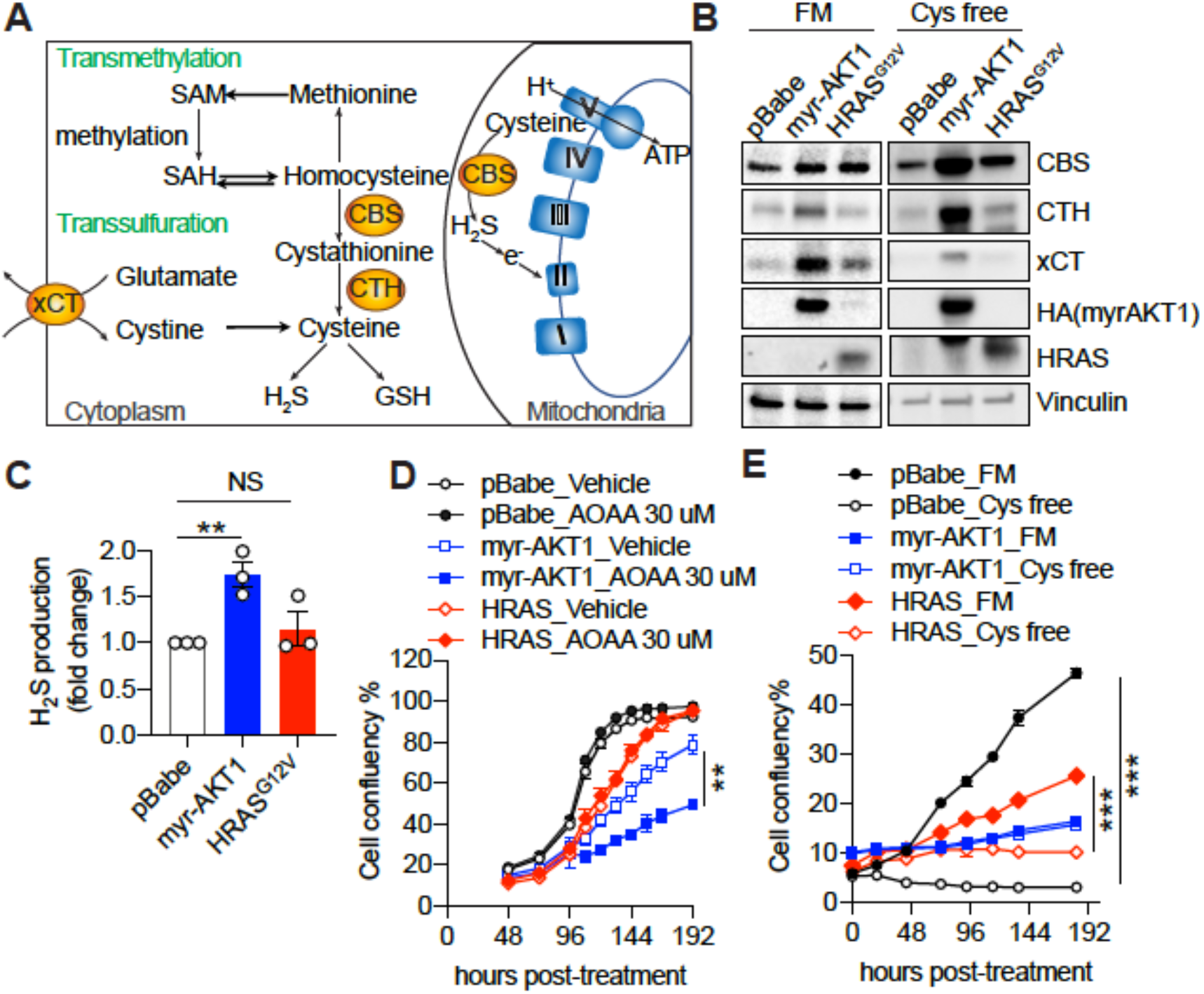
Increase of CBS expression and transsulfuration pathway activity in AKT-induced senescence. (**A**) Schematic diagram illustrating that the cytoplasmic localized CBS regulates transmethylation and transsulfuration metabolic pathways, and mitochondrial localized CBS regulates oxidative phosphorylation. (**B**) BJ-TERT cells were transduced with pBabe, myrAKT1 or HRAS^G12V^. On day 6 post-transduction the cells were plated in either full culture medium containing 100 uM cysteine (FM) or cysteine-deficient medium (Cys free). Western blot analysis was performed on day 10 post-transduction. Representative of n=3 experiments. Vinculin was probed as a loading control. (**C**) H2S production was measured by AzMC on day 14 post-transduction. Data were expressed as mean ± SEM (n = 3). **, *P* < 0.01 by one-way ANOVA. (**D**) Cells were treated with AOAA 30μM on day 5 post-transduction and cell confluency was measured by IncuCyte. Data were expressed as mean ± SEM (n = 3). (**E**) Cells were cultured in the conditions as described in (B). Cell confluency was measured by IncuCyte. Data are expressed as mean ± SEM (n = 3).

Aberrant CBS expression and/or activity contributes to a wide range of diseases including hyperhomocysteinemia (Kruger, 2017) and cancer (Zhu et al., 2018). CBS plays a complex role in cancer pathogenesis having purported tumour-promoting and -suppressive roles. Activation of CBS promoted tumour growth in colon (Phillips et al., 2017; Szabo et al., 2013), ovarian (Bhattacharyya et al., 2013), breast (Sen et al., 2015), prostate (Liu et al., 2016) and lung cancers (Szczesny et al., 2016), whereas loss of CBS in glioma cells increased tumour volume *in vivo* (Takano et al., 2014). In addition, the function of CBS in liver cancer remains inconclusive with conflicting reports of both tumour-promoting (Jia et al., 2017; Yin et al., 2012) and -suppressive roles (Kim et al., 2009). These studies underscore the context-dependent roles of CBS in cancer development.

In this study we explored the molecular mechanisms underpinning CBS’s role in maintaining AIS and how the loss of CBS promotes AIS escape. The requirement of CBS for the maintenance of AIS implicates it as a putative tumour suppressor during PI3K/AKT pathway-driven tumorigenesis. To gain insight into this, we further characterised the expression level of CBS in gastric cancer tissue samples and cells and sought to define the functional significance of CBS loss in the context of activated PI3K/AKT signalling-driven gastric cancer development.

## Results

### CBS expression level and transsulfuration pathway activity are elevated in AKT-induced senescence (AIS)

To investigate the mechanisms by which CBS contributes to AIS maintenance, we first evaluated CBS expression and activity in several non-transformed cells with hyperactivated AKT. An increase of CBS protein expression was observed in BJ-telomerase reverse transcriptase (TERT) human skin fibroblasts (Fig.1B) and IMR90 human foetal lung fibroblasts (Fig.S1A) overexpressing myristoylated (myr)-AKT1. In BJ-TERT and human mammary epithelial cells (HMEC), overexpressing AKT1^E17K^, a clinically relevant activated mutant form of AKT1 in multiple cancer types including breast cancer and ovarian cancer (Carpten et al., 2007), also enhanced CBS protein expression (Fig.S1B). However, AKT hyperactivation did not affect CBS mRNA expression (Fig S1C), suggesting a post-transcriptional regulatory mechanism underpinning increased CBS protein expression.

We hypothesised that the increased CBS expression in AKT hyperactivated cells was associated with upregulation of transsulfuration pathway activity and cysteine metabolism. We thus examined the expression levels of CTH, a key enzyme in the transsulfuration pathway and xCT, the Xc- amino acid antiporter responsible for the up-take of cystine (an oxidised form of cysteine) (Fig 1A). Both CTH and xCT were upregulated during AIS compared to proliferating control cells, suggesting an elevated cysteine synthesis via the transsulfuration pathway and cysteine uptake (Fig. 1B). In contrast, senescent cells expressing HRAS^V12^ cells exhibited a moderate increase of CBS but no change in CTH expression level. An increase in xCT was also observed during RIS, albeit to a lesser extent than during AIS, in line with the finding that upregulation of xCT facilitates RAS-mediated transformation (Lim et al., 2019).

To assess transsulfuration pathway activity, we measured H2S production. A significant increase in transsulfuration pathway activity was observed in BJ-TERT fibroblasts upon AKT but not HRAS hyperactivation (Fig.1C), suggesting that activation of transsulfuration pathway is a specific cellular response to constitutive activation of AKT. Cysteine starvation has been reported to induce necrosis and ferroptosis in cancer cells (Chen et al., 2017). Since the transsulfuration pathway mediates *De Novo* cysteine synthesis, an increase in transsulfuration pathway activity may support survival of AKT expressing cells upon cysteine limitation. Consistent with this hypothesis, inhibition of H_2_S production by aminoxyacetate (AOAA) (Szabo, 2016) selectively impaired cell proliferation (Fig. 1D) and increased SA-βGal activity (Fig. S1D) of BJ-TERT cells overexpressing myr-AKT1, suggesting that the transsulfuration pathway has a protective effect on cells expressing hyperactivated AKT. Furthermore, cysteine deprivation potently increased the expression levels of CBS and CTH in AIS cells (Fig.1B) and did not affect the survival of AIS cells, consistent with increased cysteine synthesis due to elevated CBS expression being critical for cell viability (Fig.1E).

### Depletion of CBS promotes escape from AKT-induced senescence

To validate the functional role of CBS in AIS maintenance, we depleted CBS using two independent shRNAs in BJ-TERT cells overexpressing myr-AKT1 (Fig.2A). Loss of CBS in AIS cells significantly decreased the proportion of cells with SA-βGal activity, increased EdU incorporation and enhanced colony formation, demonstrating an essential role of CBS in AIS maintenance (Fig 2B). To further confirm the on-target specificity of the knockdown, we generated BJ-TERT cells expressing a doxycycline-inducible CBS shRNA and an shRNA-resistant 4-OHT-inducible estrogen receptor (ER)-tagged-CBS fusion (Fig.2C and S2A). Upon expressing myr-AKT1, these cells underwent AIS, as indicated by a significant increase in SA-βGal-positive cells and decrease in EdU-positive cells and, consistent with the finding in the siRNA AIS escape screen. CBS depletion induced by doxycycline decreased SA-βGal activity and increased EdU incorporation in AIS cells (Fig.2D). Importantly, simultaneously depleting CBS and expressing ER-CBS prevented senescence escape, confirming the on-target specificity of the knockdown and the critical role of CBS in maintaining AIS (Fig.2D). Modulation of CBS expression in proliferating (pBabe) control cells did not affect the percentage of SA-βGal- and EdU-positive cells (Fig.2D), suggesting that CBS is a downstream effector of AKT and its depletion alone is insufficient to induce senescence. Similar to the findings in BJ-TERT cells, AKT1 hyperactivation also caused senescence in IMR-90 lung fibroblasts (Fig S2B). CBS knockdown in AIS cells significantly suppressed SA-βGal staining and enhanced EdU incorporation but not in proliferating cells (Fig S2B). Knockdown of CBS in BJ-TERT cells with constitutive RAS activation did not affect colony formation, suggesting CBS has a specific regulatory role during AIS but not RIS maintenance (Fig.2E and S2C).

**Figure 2.**
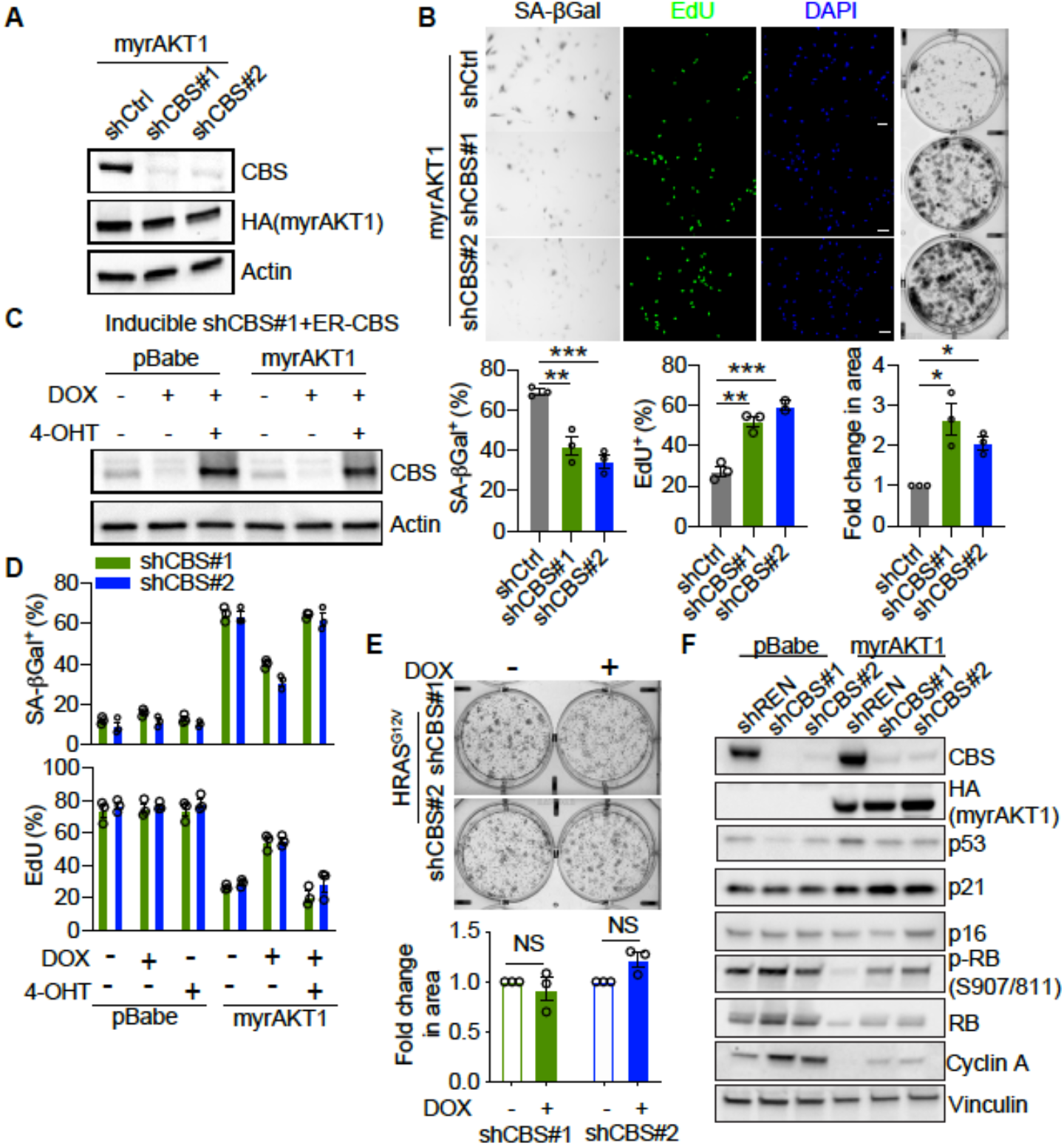
Depletion of CBS promotes escape from AKT-induced senescence. (A and B) BJ-TERT cells transduced with myrAKT1 were transduced with pGIPZ-shCBS or control vector pGIPZ-NTC (shCtrl) on day 6 post-transduction of myrAKT1. (**A**) Western blot analysis was performed on day 8 post-transduction of shRNA. Representative of n = 2 experiments. (**B**) Images and quantification of the percentage of cells with positive staining for SA-βGal activity and EdU incorporation on day 8 post-transduction of shRNA, as well as colony formation assay on day 14 post-transduction of shRNA. Data were expressed as mean ± SEM (n = 3). *, *P* < 0.05; **, *P* < 0.01; ***, *P* < 0.001 by one-way ANOVA. (**C-E**) BJ-TERT cells expressing doxycycline-inducible CBS shRNA#1 and 4-OHT-inducible CBS were transduced with pBabe or myr-AKT1, treated with doxycycline (1μg/ml) ± 4-OHT (20 nM) on day 5 post-transduction and analyzed on day 14 post-transduction. (**C**) Western blot analysis of CBS expression. Actin was probed as a loading control. (**D**) Quantification of the percentage of cells with positive staining for SA-βGal activity or EdU proliferation marker incorporation. Data were expressed as mean ± SEM (n = 3). (**E**) BJ-TERT cells expressing doxycycline-inducible CBS shRNA were transduced with HRAS^G12V^, treated with doxycycline (1μg/ml) on day 5 post-transduction and colony formation assay analyzed on day 14 post-transduction. Data were expressed as mean ± SEM. n = 3 experiments. NS, not significant by student’s t test. (**F**) BJ-TERT cells expressing doxycycline-inducible shCBS or control shREN were transduced with pBabe or myr-AKT1 and then treated with doxycycline (1μg/ml) on day 5 post-transduction. Western blot analysis of CBS, AKT and the key senescence markers as indicated was performed on day 14 post-transduction. Vinculin was probed as a loading control. Representative of n=2 experiments.

To determine the mechanisms by which CBS depletion causes escape, we examined the impact on additional senescence hallmarks. While loss of CBS released AIS cells from cell cycle arrest, knockdown of CBS did not significantly change the mRNA and protein expression level of several key SASP-related genes including IL-1α, IL-1β, IL-6 and IL-8, which are upregulated during AIS (Astle et al., 2012; Chan et al., 2020) (Fig. S2D and S2E). Given the p53 and Retinoblastoma protein (Rb) pathways predominantly control senescence-mediated cell cycle arrest, we examined signalling downstream of AKT activation in the presence and absence of CBS (Fig 2F). Consistent with our previous findings, p53 and its downstream target p21 were upregulated during AIS, but depletion of CBS had no effect on these levels. Furthermore, total Rb, a key regulator of the G1/S phase transition, and its inhibitory phosphorylated form at serine 807/811 were markedly suppressed in AIS and partially rescued upon CBS knockdown. Cyclin A, which mediates S to G2/M phase cell cycle progression, was also upregulated upon depleting CBS in cells with hyperactivated AKT. These results demonstrate that CBS depletion can restore the proliferation of cells that have undergone AIS in a p53-independent manner.

### Depletion of CBS in AIS cells increases GSH metabolism in cysteine-replete conditions

CBS is the key enzyme regulating transsulfuration and transmethylation pathways. By analysis of the data from the AIS-escape siRNA screen, we found that except CBS, siRNA knockdown of other genes involved in transsulfuration and transmethylation pathway did not significantly affect AIS cell numbers (robust Z score < 2, Table S4). Therefore, it is likely that AIS escape in cysteine-replete conditions by loss of CBS is through a transsulfuration/transmethylation pathway-independent mechanism. In addition, knockdown of CBS did not affect the expression levels of CTH and xCT when BJ-TERT cells were grown in cysteine-replete culture medium (Fig.S3A).

To further determine the activities of these two metabolic pathways in AIS cells and the effect of CBS depletion, we assessed the thiol redox metabolome using LC/MS after thiol derivatisation with N-Ethylmaleimide (NEM) (Fig.S3A-I and Table S5). Compared to proliferating cells (pBabe-siOTP), AIS cells (myr-AKT1-siOTP) exhibited increased abundance of homocysteine (hcy), methionine, SAM and SAH (Fig.3A-D), indicating upregulation of transmethylation pathway activity in AIS. Meanwhile, the level of cysteine was also increased (Fig.3F) whereas the level of cystathionine decreased (Fig.3E) in AIS cells, supporting that AKT activation stimulates the transsulfuration pathway, consistent with the increased H2S production capacity in AIS cells (Fig. 1C). Activation of the transsulfuration pathway can also increase GSH synthesis. Indeed, AIS cells exhibited a much higher level of the GSH synthesis precursor γ-Glu-Cys (Fig.3H) and the GSH degradation product Cys-Gly (Fig.3I) than proliferating cells despite lack of change in GSH (Fig.3G), suggesting an elevated GSH metabolism in AIS cells.

**Figure 3.**
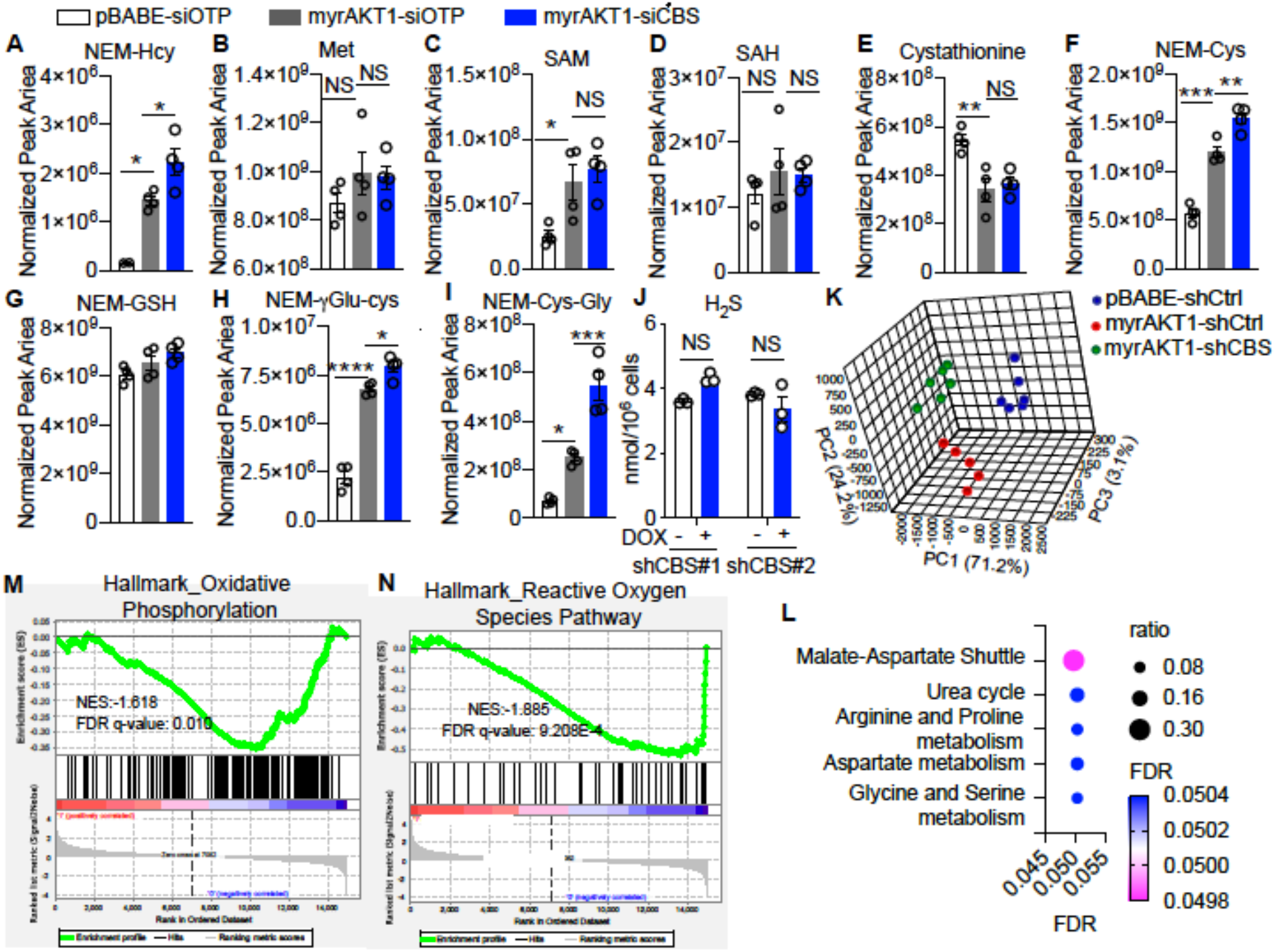
Depletion of CBS in AIS cells does not alter activities of transmethylation and the transsulfuration pathways. (**A-I**) BJ-TERT cells were transduced with pBabe or myrAKT1. After four days cells were then transfected with either CBS siRNA (siCBS) or control siRNA (siOTP) and analyzed on day 6 post-siRNA transfection. The thiol redox metabolome was assessed by targeted LC/MS. The abundance of metabolites including (**A**) NEM-Hcy, (**B**) Methionine (Met), (**C**) SAM, (**D**) SAH, (**E**) cystathionine, (**F**) NEM-Cys, (**G**) NEM-GSH, (**H**) NEM-γ-Glu-Cys and (**I**) NEM-Cys-Gly were shown. Data were expressed as mean ± SEM (n = 4). (**J**) BJ-TERT cells expressing doxycycline-inducible shCBS were transduced with myrAKT1, and treated with doxycycline (1μg/ml) on day 5 post-transduction. H2S production capacity was measured by AzMC on day 14 post-transduction. Data are expressed as mean ± SEM (n = 3). *, *P* < 0.05; **, *P* < 0.01; ****, *P* < 0.0001 by one-way *ANOVA*. (**K**) BJ-TERT cells were transduced with pBabe or myr-AKT1 followed by transduction with pGIPZ-shCBS (myrAKT1-shCBS) or control vector pGIPZ-NTC (pBABE-shCtrl, myrAKT1-shCtrl) on day 6 post-transduction. The metabolic profiling by GC/MS was performed on day 14 post-transduction. n = 6 in both pBabe-shCtrl and myrAKT1-shCBS groups and n = 5 in myrAKT1-shCtrl group, with one sample excluded due to a technical issue in sample processing. PCA plot, x, y and z-axis represent individual principle component. (**L**) Metabolite Sets Enrichment Analysis of myrAKT1-shCBS comparing with myrAKT1-shCtrl cells. Ratio is the value obtained by dividing the number of metabolites in our data assigned to a metabolic process with the number of metabolites curated in the same process in MetaboAnalyst 4.0. (**M and N**) Gene set enrichment analysis of RNA-seq data showing downregulation of hallmark of oxidative phosphorylation and reactive oxygen species pathways in myrAKT1-shCBS cells compared with myrAKT1-shCtrl cells.

Loss of CBS can cause upstream substrate hcy accumulation. Indeed, compared to AIS cells with intact CBS, an elevated hcy level in CBS-depleted cells (myrAKT1-siCBS) was detected (Fig.3A). However, the metabolites in the transmethylation pathway including methionine (Fig.3B), SAM (Fig. 3C) and SAH (Fig. 3D) were unaffected by CBS knockdown. In terms of transsulfuration pathway, knockdown of CBS, unexpectedly, further increased the abundance of the metabolites including cysteine (Fig.3F), γ-Glu-Cys (Fig. 3H) and Cys-Gly (Fig.3I), suggesting loss of CBS further stimulates GSH metabolism. In contrast, H2S production was not affected by CBS knockdown (Fig.3J). Upregulation of GSH metabolism upon CBS depletion implicated that senescence escape due to CBS knockdown is associated with regulation of oxidative stress and antioxidation activity.

To explore whether other metabolic alterations are associated with CBS-mediated AIS maintenance, we performed gas chromatography mass spectrometry (GC/MS)-based untargeted metabolomics (Table S6 and S7). BJ-T cells transfected with a control vector (proliferating), myr-AKT1 (AIS) and myr-AKT1+shCBS (AIS escaped) displayed distinct metabolic profiles, as indicated by Principle Component Analysis (PCA) (Fig.3K). Metabolic pathway enrichment analysis revealed that the top pathway re-wired in CBS-depleted cells compared to AIS cells was the malate-aspartate shuttle with increase of glutamic acid and aspartic acid (Fig.3K. Fig.S3B and Table S8), which is the key mechanism for transporting NADH from the cytoplasm to the mitochondria in order to support oxidative phosphorylation. Thus, alteration of mitochondrial function required to maintain AIS may be associated with AIS escape caused by CBS depletion.

To further investigate the molecular mechanisms of how CBS maintains AIS, we characterised the transcriptomic changes upon depleting CBS during AIS. RNA-seq was performed on BJ-TERT cells expressing myrAKT1 transduced with control or CBS shRNA. Differential gene expression analysis of CBS-depleted AIS cells compared with control AIS cells revealed 404 genes were significantly up-regulated (adjusted p-value < 0.05, Log_2_FC > 1) and 181 genes significantly down-regulated (adjusted p-value < 0.05, Log_2_FC < −1) (Fig.S3C and Table S9 and S10). Gene set enrichment analysis (GSEA) using the hallmark gene sets in the Molecular Signatures Database (MSigDB) identified that pathways involved in oxidative phosphorylation and reactive oxygen species (ROS) were significantly downregulated in CBS-depleted AIS cells compared to the control AIS cells (Fig.3L and 3M, Table S11 and S12). Collectively, these data implicate altered mitochondrial energy metabolism and ROS production contribute to CBS-dependent AIS maintenance.

### CBS deficiency alleviates oxidative stress in AIS cells

CBS has been reported to localise to both the cytoplasm and mitochondria and regulate mitochondrial function and ATP synthesis via H2S (Bhattacharyya et al., 2013; Panagaki et al., 2019). Consistent with this, we also observed CBS localisation in the mitochondria (Fig.4A) by immunofluorescent cell staining with mitoTracker. AIS cells exhibited elevated mitochondrial abundance as indicated by increased intensity of mitoTracker staining compared to the proliferating cells (Fig.4A) and an increased abundance of proteins involved in the mitochondrial electron transport chain (Fig.S4A).

**Figure 4.**
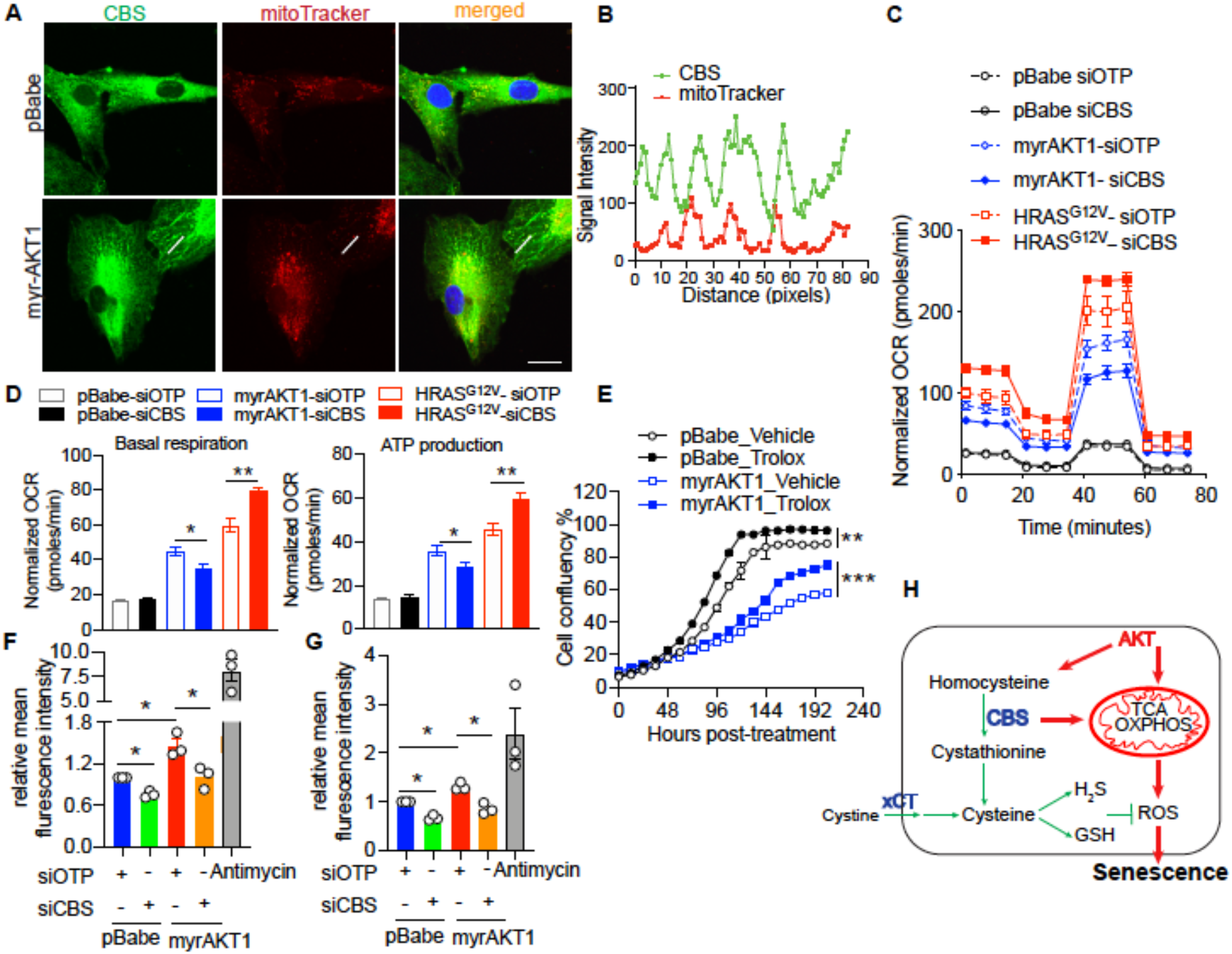
CBS deficiency alleviates oxidative stress in AKT-induced senescent cells. (**A-B**) BJ-TERT cells were transduced with pBabe, myrAKT1. Immunofluorescent staining showing CBS (Green) and mitochondria (Red) on day 10 post-transduction. The representative images are from one of two independent experiments. (**B**) Quantification of signal intensities using ImageJ by applying a single ROI to two color channels in the same image and extracting the plot profile. Scale bar = 20 μm. (**C-D**) BJ-TERT cells were transduced with either pBabe, myrAKT1 or HRAS^G12V^. After five days cells were then transfected with either CBS siRNA (siCBS) or control siRNA (siOTP) and analyzed on day 6 post-siRNA transfection. (**C**) Oxygen consumption rate (OCR) was determined using the Seahorse XF96 Extracellular Flux Analyzer. A mitochondrial stress test was carried out by sequential injection of oligomycin (1μM), FCCP (1μM) and rotenone/antimycin (0.5 μM each). Data were expressed as mean ± SEM of 4 technical replicates. N = 2. (**D**) The basal respiration rate and the ATP production rate determined by the mitochondrial stress test. *, *P* < 0.05; **, *P* < 0.01 by student’s t test. (**E**) BJ-TERT cells were transduced with either pBabe or myrAKT1. On day 5 post-transduction, cells were treated with Trolox 100uM. Cell confluency was measured by IncuCyte. Data were expressed as mean ± SEM (n = 3). (**F and G**) Flow cytometric analysis of the mitochondrial superoxide production by MitoSOX (**F**) and the cytoplasmic ROS production by H2DCFDH-DA on day 6 post-siRNA transfection (**G**). Data are expressed as mean ± SEM (n = 3). *, *P* < 0.05 by one-way ANOVA. (**H**) Schematic diagram illustrating CBS-mediated metabolic alterations on maintenance of AKT-induced senescence (AIS). CBS stimulates mitochondrial oxidative phosphorylation and consequently increases ROS production which contributes to AIS maintenance. On the other hand, AKT hyperactivation stimulates transsulfuration pathway activity with consequent enhanced H2S and GSH production which protects cells from excessive oxidative stress-induced cell death.

To investigate the role of CBS-mediated mitochondrial alterations in AIS maintenance, we examined the oxidative phosphorylation status in AIS cells transfected with control or CBS siRNA by measuring the oxygen consumption rate (OCR) using a Seahorse extracellular flux analyser. OCR was markedly elevated during AIS (Fig.4C) and knockdown of CBS significantly suppressed basal OCR and ATP production (Fig.4D). These results suggest that CBS is required for enhanced oxidative phosphorylation during AIS maintenance and CBS depletion reduces mitochondrial bioenergetics in AIS. To test whether these effects were specific for AIS, we also performed Seahorse analysis on cells during RIS. While basal OCR was also increased during RIS, knockdown CBS in RIS cells, in contrast to those undergoing AIS, had an opposite effect on oxidative phosphorylation by further upregulating basal OCR and ATP production (Fig.4D). This distinct effect on mitochondrial activity is likely to contribute to the specific regulatory role of CBS in AIS maintenance.

Mitochondria are the major intracellular organelles of ROS production. Elevated ROS results in oxidative stress which may underlie AKT-induced senescence. To test this hypothesis, we first treated AIS cells with an antioxidant Trolox, which resulted in increased proliferation (Fig. 4E) and decreased SA-βGal staining (Fig. S4B), establishing the role of oxidative stress in AIS maintenance. To test the impact of AKT hyperactivation on mitochondrial and cytoplasmic ROS production, we performed flow cytometry analysis using MitoSOX and H2DCFDH-DA, respectively, on proliferating and AIS cells transfected with control or CBS siRNA. Critically, CBS knockdown decreased ROS levels in AIS cells (Fig.4F and 4G and Fig.S4C). Together, these results strongly support the concept that increased oxidative phosphorylation and ROS production sustain AIS status and are impaired by CBS loss. In parallel, AKT activation increases transsulfuration pathway activity, which consequently stimulates GSH and H2S production, thereby protecting AIS cells from ROS-induced cell death. Notably, the antioxidant activity of this metabolic pathway is still retained and GSH metabolism is even further enhanced upon CBS loss. Together the alleviated oxidative stress contributes to escape of AIS cells from cell cycle arrest (Fig. 4H).

### CBS expression is frequently suppressed in gastric cancer

Given that we showed CBS loss promotes escape from AIS, we hypothesised that loss of CBS could cooperate with oncogenic activation of the PI3K/AKT/mTORC1 pathway to promote tumourigenesis. Analysis of TCGA stomach adenocarcinoma data from 478 samples using cBioPortal (http://www.cbioportal.org) identified CBS deep deletions and mutations in gastric cancer (Fig.5A). Further analysis of the TCGA gastric cancer patient data revealed that PI3K/AKT/mTORC1 signalling pathway alterations occur in 33% of gastric cancer with PIK3CA (15%) and PTEN (9%) alterations being the most common genetic alterations (Fig.S5A). CBS genetic alterations were found in 3% of gastric cancers including 4 cases with CBS deep deletions co-occurring with hyperactive PI3K/AKT/mTORC1 signalling (Fig.S5A). Furthermore, CBS mRNA level was significantly decreased in tumours (N=406) compared to adjacent normal tissues (N=211) in gastric cancer patients (Fig.5B), consistent with the hypothesis that CBS is a tumour suppressor in gastric cancers characterised by PI3K/AKT/mTORC1 pathway alterations.

**Figure 5.**
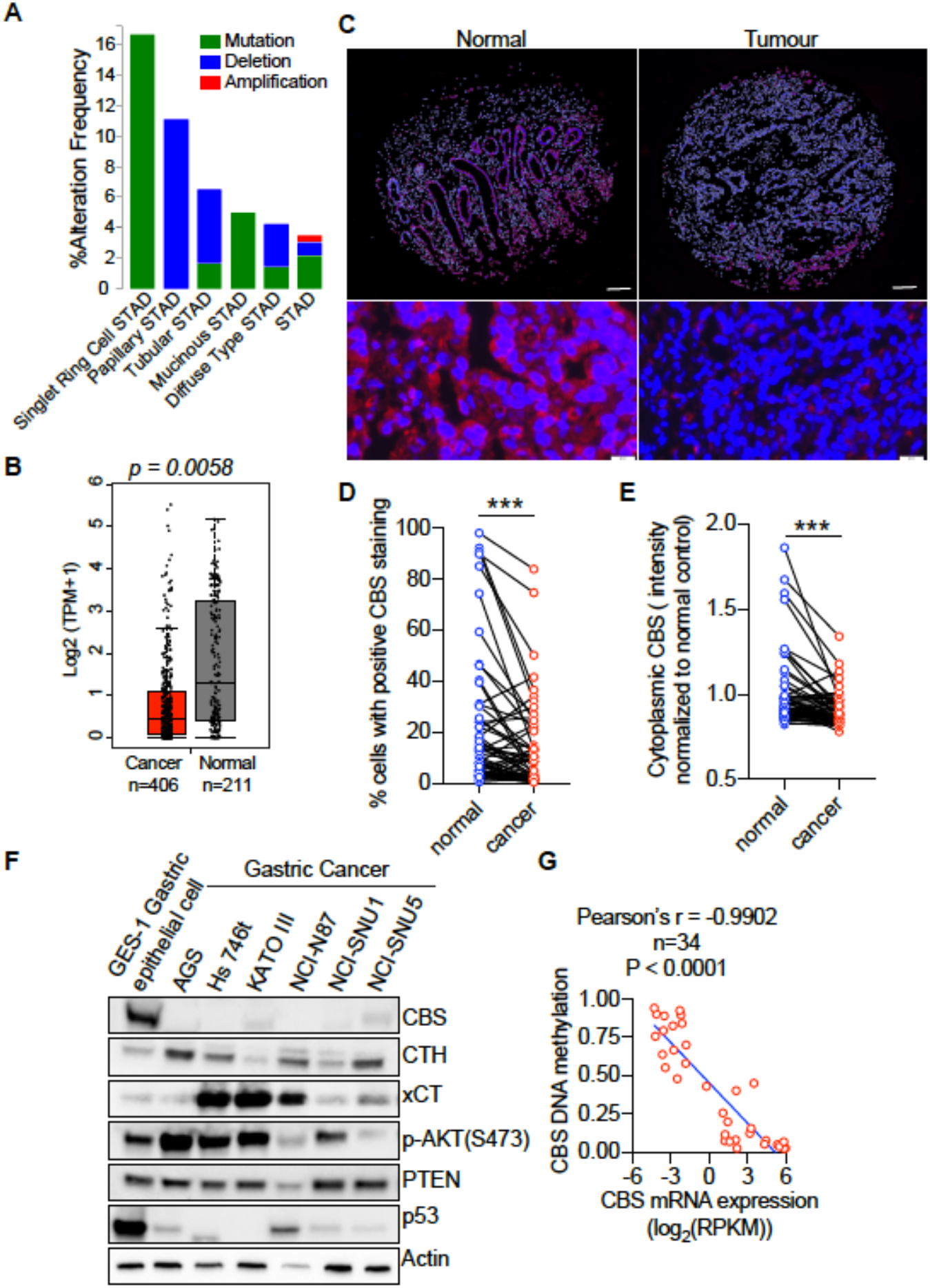
CBS expression is suppressed in tumor tissues and human cell lines of gastric cancer. (**A**) Analysis of genetic alterations of CBS in different subtypes of gastric cancer using the cBioPortal (http://www.cbioportal.org). (**B**) Median mRNA expression level of CBS in normal stomach tissues and stomach adenocarcinoma tissues profiled by Gene Expression Profiling Interactive Analysis (GEPIA, http://gepia.cancer-pku.cn) based on the TCGA database. *P* = 0.0058. (**C-E**) Gastric cancer patient tissue microarray was assessed by immunofluorescent staining of CBS (red) and counterstained for the nucleus (DAPI, blue). (**C**)The representative images of normal or cancer tissues from one gastric cancer patient were shown, Top panel: Scale bars = 200 μm. Bottom panel: Scale bars = 50 μm. The percentage of cytosolic CBS-positive cells (**D**) and the intensity of cytoplasmic CBS (**E**) in the tumor or adjacent normal tissues from 62 patients were shown. ***, *P* < 0.001 by paired student’s t-test. (**F**) Western blot analysis of cysteine metabolism-associated proteins and PTEN-AKT signaling in GES-1 gastric epithelial cell line and six gastric cancer cell lines. Actin was probed as a loading control. Representative of n = 3 experiments. (**G**) Correlation of CBS mRNA expression with CBS DNA methylation in 34 gastric cancer cell lines based on the data retrieved from the Cancer Cell Line Encyclopedia.

To evaluate alteration of CBS protein expression in human gastric cancer, we assessed CBS protein levels in paired samples of gastric tumours and adjacent non-cancerous mucosa from 62 gastric cancer patients in a tissue microarray using immunofluorescent staining (Fig.5C). This tissue microarray was assembled from paraffin-embedded tissue blocks collected from gastric cancer patients who underwent gastrectomy from 2000 to 2005 in Changhai Hospital, Shanghai, China as described previously (Zhang et al., 2013). Cytosolic CBS protein expression level, as measured by the percentage of cells with positive CBS staining (Fig.5D), and fluorescence intensity (Fig.5E) were significantly down-regulated in tumour tissues compared to the adjacent normal gastric tissues.

To establish a cell-based system to probe the interaction between activated PI3K/AKT/mTORC1 signalling and loss of CBS expression, we first examined CBS protein expression in six gastric cancer cell lines compared with a SV40-transformed gastric epithelial cell line GES-1, which was derived from foetal stomach mucosa and was non-tumourigenic in nude mice (Ke et al., 1994). Compared to gastric epithelial cells, CBS expression was markedly decreased in all gastric cancer cell lines while elevated AKT activity, as indicated by increase of AKT phosphorylation, was observed in in AGS, Hs746T, KATO III gastric cancer cell lines (Fig.5F).

Previous studies have demonstrated CBS deficiency in human gastric cancer cells with hyper-methylation at the CBS promoter region, implicating loss of CBS through epigenetic regulation in gastric cancer (Zhao et al., 2012). Comparison of CBS mRNA expression with DNA methylation status in a panel of 34 human gastric cancer cell lines revealed a significant negative correlation between CBS mRNA expression level and DNA methylation (Fig.5G). Indeed, the CBS promoter was methylated in all six gastric cancer cell lines tested, and half of them displayed complete absence of unmethylated CBS promoter (SNU-1, NCI-N87 and AGS) that was associated with undetectable CBS transcription (Fig.S5D). Blocking DNA methylation with Azacitidine, a DNA methyltransferase (DNMT)-inhibiting cytosine nucleoside analogue, up-regulated CBS mRNA expression in all cell lines tested except Hs746T, whereby SNU-1 cells showed a >50-fold increase, confirming epigenetic silencing of CBS expression in gastric cancer cells (Fig.S5E). Given the GES-1 cells express high levels of CBS as well as activated AKT signalling comparable to some of gastric cancer cell lines, we used this cell line to interrogate the role of CBS loss in the initiation of gastric cancer.

### Loss of CBS cooperates with PI3K/AKT pathway activation to promote gastric cancer pathogenesis

To further test if CBS loss cooperates with PI3K/AKT/mTORC1 hyperactivation in gastric cancer oncogenesis, we transduced myrAKT1 and CBS/control shRNA into GES-1 cells and tested their ability to form colonies in soft agar (Fig.6A). Consistent with our results in fibroblasts, AKT hyperactivation increased transsulfuration pathway activity and GSH production, which was not affected by CBS depletion (Fig. S6A and S6B). Hyperactivation of AKT1 had no significant effect on cell proliferation in 2D culture and did not induce senescence due to inactivation of p53 and pRb by SV40T in the GES-1 cells (Fig. S6C). However, it effectively promoted anchorage-independent growth, and this was further enhanced by CBS depletion (Fig.6A and 6B). CBS depletion also significantly enhanced anchorage-independent growth of GES-1 cells with clustered regularly interspaced short palindromic repeats (CRISPR)/Cas9-mediated PTEN knockout (Fig.6C and 6D) but not in 2D cell culture (Fig. S6D). These results support that CBS loss can cooperate with PI3K/AKT signalling to promote oncogenic transformation.

**Figure 6.**
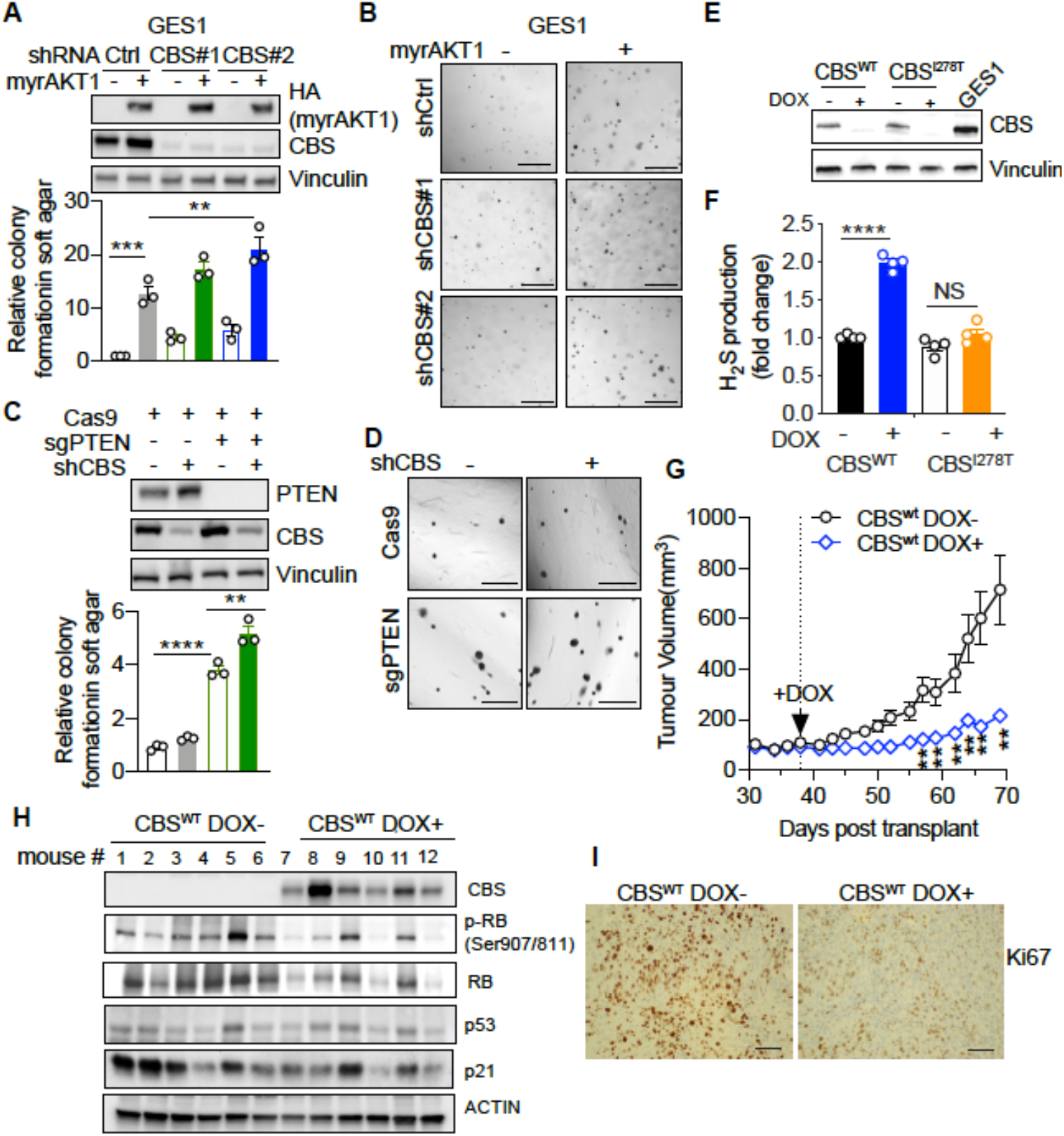
Loss of CBS synergizes with PI3K/AKT pathway to promote gastric cancer pathogenesis. (**A-B**) GES-1 gastric epithelial cells were stably transfected with doxycycline-inducible myrAKT1 and pGIPZ-shCBS or control pGIPZ-NTC (shCtrl). (**A**) Top: Western blot analysis showing HA-tagged myrAKT1 and CBS after doxycycline (0.75 μg/ml) induction for 3 days. Vinculin was probed as a loading control. Bottom: The anchorage-independent growth was assessed by soft agar colony formation assay. The number of colonies formed were counted on day 21 post-doxycycline (0.75 μg/ml) induction. Data were expressed as mean ± SEM (n = 3). **, *P* < 0.01, ***, *P* < 0.001, by one-way ANOVA. (**B**) The representative images of colonies formed in the soft agar. Scale bar = 500 μm. (C-D) GES-1 cells with PTEN knockout by CRISPR or Cas9 control were transduced with doxycycline-inducible CBS shRNA#1. (**C**) Top: Western blotting showing PTEN and CBS expression after treatment with doxycycline (0.75 μg/ml) for 3 days. Vinculin was probed as a loading control. Bottom: The anchorage-independent growth was assessed by soft agar colony formation assay after doxycycline induction for 21 days. Data were expressed as mean ± SEM (n = 3). **, *P* < 0.01; ****, *P* < 0.0001 by one-way ANOVA. (**D**) The representative images of colonies formed in the soft agar. Scale bar = 500 μm. (**E-F**) AGS gastric cancer cells were stably transfected with doxycycline-inducible CBS^wt^ or CBS^I278T^. Cells were cultured in the presence or absence of doxycycline induction (0.08 μg/ml for CBS^wt^ or 1 μg/ml for CBS^I278T^). (**E**) Western blot analysis on day 3 post-treatment. Vinculin was probed as a loading control. (**F**) H2S production. Data are expressed as mean ± SEM. n = 4 experiments. (G-I) AGS gastric cancer cells expressing doxycycline-inducible CBS^wt^ were subcutaneously implanted in BALB/c nude mice. On day 38 post-implantation, mice were supplied with water containing 0.2% (W/V) doxycycline and 600 mg Doxycycline/kg food. (**G**) Tumor volume measured in AGS xenografts. Data are expressed as mean ± SEM. n=6 mice per group. (**H**) Western blot analysis of CBS and senescence-associated protein expression in tumor tissues. Actin was probed as a loading control. (**I**) IHC of Ki67 on the tumor tissue. Scale bar: 50 μm.

To further investigate the functional cooperation of CBS and PI3K/AKT signalling in gastric cancer pathogenesis, we engineered AGS gastric cancer cells, which harbour *CBS* deficiency and *PIK3CA* mutations E545A and E453K resulting in AKT activation (Fig.5F), to express a doxycycline-inducible wild type CBS or an inactive CBS^I278T^ mutant. This mutation is the most frequently observed CBS mutation in cancer cells and exhibits only ~2.4% of the enzyme activity of wild type CBS (Kruger & Cox, 1995). Treatment of AGS cells with doxycycline restored CBS^wt^ and CBS^I278T^ protein expression to a level comparable to that of GES-1 gastric epithelial cells (Fig.6E). Restoration of wild type CBS increased H2S production (Fig.6F) but did not significantly enhance GSH abundance (Fig.S6E). Interestingly, restoration of CBS expression did not affect AGS cell proliferation (Fig.S6F).

To evaluate the functional impact of CBS restoration *in vivo*, we transplanted the AGS cells expressing doxycycline-inducible CBS into immunocompromised mice. Induction of CBS^wt^ significantly suppressed AGS tumour growth (Fig.6G). Induction of CBS was also associated with a marked decrease in Ki67 expression and inhibitory RB phosphorylation without altering p53 and p21 expression levels in the tumour tissues, suggesting that restoration of CBS expression could suppress gastric tumour formation independent of p53 (Fig.6H and 6I).

## Discussion

Hyperactivation of the PI3K/AKT/mTORC1 signaling pathway causes a senescence-like phenotype in non-transformed cells, which acts as a protective brake against tumor formation (Zhu et al., 2020). Subsequent genetic or epigenetic changes can disengage this brake and lead to oncogenic transformation. Deregulated metabolism along with cell cycle withdrawal, SASP and macromolecular damage, are hallmarks of the senescence phenotype (Gorgoulis et al., 2019). Oxidative stress is a key metabolic feature of RAS-induced senescence and ROS triggers DNA damage and proliferative arrest in RIS cells (Irani et al., 1997; Lee et al., 1999; Ogrunc et al., 2014). Activation of AKT can also increase intracellular ROS levels by stimulating oxidative phosphorylation and impairing ROS scavenging by inhibition of FoxO transcription factors (Nogueira et al., 2008). Consistent with these findings, we demonstrate increased mitochondrial abundance and respiratory activity, as well as ROS production in the senescent-like cells resulting from AKT hyperactivation. Alleviation of oxidative stress by antioxidant treatment partially releases AIS cells from the state of cell division arrest, supporting that ROS is required for AIS maintenance.

On the other hand, activation of the PI3K/AKT pathway has been observed to induce a potent antioxidant response (Hoxhaj & Manning, 2020) that may antagonise the tumour-suppressive AIS. One major ROS-scavenging mechanism by the PI3K/AKT pathway is through sustained activation of nuclear factor erythroid 2-related factor 2 (NRF2) (Mitsuishi et al., 2012; Rada et al., 2011). In this study we uncovered another mechanism of AKT-mediated ROS detoxification by upregulation of transsulfuration pathway activity and enhancing glutathione and H2S synthesis (Fig.4H). The key metabolic enzymes in the transsulfuration pathway including CBS and CTH have been reported to be positively regulated through the PI3K/AKT/Sp1 axis, and inhibition of PI3K pathway has been shown to decrease CBS and CTH expression levels (Liu et al., 2017; Yin et al., 2012). Meanwhile, we found that AKT activation markedly increased xCT protein expression, a cystine-glutamate antiporter encoded by *SLC7A11*, indicating the increase of extracellular cystine uptake in AIS. Consequently, an elevated intracellular cysteine level promotes the production of GSH and H2S, the critical components in the antioxidant system. Previous studies have shown that H2S inhibits H_2_O_2_-mediated mitochondrial dysfunction by preserving the protein expression levels and activity of key antioxidant enzymes, inhibiting ROS production and lipid peroxidation (Wen et al., 2013). Moreover, these effects may be associated with sulfhydration of Keap1 and activation of Nrf2 or increased production of the antioxidant glutathione (Koike et al., 2013; Yang et al., 2013). In addition to antioxidant effects, H2S can modulate mitochondrial functions and cellular bioenergetics in a concentration-dependent manner. At low concentrations, H2S acts as a mitochondrial electron donor to mitochondrial complex II, resulting in bioenergetic stimulation (Szabo et al., 2013). At high concentrations, H2S acts as a mitochondrial poison via the inhibition of cytochrome c oxidase in mitochondrial complex IV (Panagaki et al., 2019; Szabo et al., 2014). Our finding that suppression of H2S by AOAA exacerbates proliferative arrest of AKT-hyperactivated cells supports a protective effect of H2S from oxidative stress-induced cell death (Fig.1D). The protective role of the transsulfuration pathway in AIS is further supported by the finding that myrAKT1-expressing cells are resistant to exogenous cysteine deprivation (Fig. 1E). Thus, we propose that increased levels of GSH and H2S through the transsulfuration pathway during AIS maintenance enhance the antioxidant capacity of AIS cells, protecting senescent cells from ROS-induced cell death (Fig.4H).

Startlingly, we found AIS cells escape upon CBS depletion under cysteine-replete conditions. This is mediated by decrease of ROS production through suppression of mitochondrial oxidative phosphorylation and increase of GSH metabolism (Fig. 4H). Intriguingly, a recent publication reported that overproduction of H2S by increased mitochondrial-localised CBS expression results in suppression of mitochondrial oxidative phosphorylation and ATP production in the fibroblasts from Down syndrome patients (Panagaki et al., 2019). This discrepancy could be partially explained by the bell shaped or biphasic biological effect of H2S as mentioned above. It is unclear how loss of CBS increased GSH metabolism. One explanation is that loss of CBS reciprocally upregulates xCT activity and leads to a compensatory upregulation of cystine uptake and GSH synthesis.

Oncogene-induced senescence acts as a critical tumor-suppressive brake and this senescence brake is disengaged during tumorigenesis. Based on our observation that loss of CBS promoted AIS escape in normal cells, we propose a potential tumour-suppressive role for CBS in cancers harbouring PI3K/AKT pathway activation. We demonstrated suppression of CBS expression in primary gastric tumour tissues and a panel of human gastric cancer cell lines through epigenetic silencing. In parallel, a negative correlation between CBS and xCT expression was observed in gastric cancer cell lines, which is consistent with a recent report that cancer cells maintain intracellular cysteine levels and sustain cell proliferation by increasing xCT-mediated cystine uptake (Zhu et al., 2019). We further demonstrated that loss of CBS cooperates with AKT hyperactivation to promote anchorage independent growth of gastric epithelial cells and restoration of CBS expression inhibited PI3K/AKT hyperactive gastric tumour growth. Induction of apoptosis and impairment of gastric cancer cell metastasis by increasing H2S production upon NaHS treatment has been previously reported (Zhang et al., 2015). Whether the tumour-suppressive effect of CBS in gastric cancer cells is mediated through H2S requires further investigation.

Taken together, our study identifies CBS as a novel regulator of AIS maintenance and a potential tumour suppressor in gastric cancer pathogenesis, potentially providing a new metabolic vulnerability that can be harnessed to target PI3K/AKT/mTORC1-driven cancers.

## Methods

### Cell culture and reagents

BJ-TERT immortalized human foreskin fibroblasts were a gift from Robert Weinberg (Massachusetts Institute of Technology, USA). Primary IMR-90 lung fibroblasts originating from the American Type Culture Centre (ATCC-CL-186) were obtained from the Garvan Institute of Medical Research, Sydney, Australia. Human embryonic kidney (HEK293T) cells were purchased from the ATCC (ATCC-CRL-3216). Human gastric cancer cell lines, AGS (ATCC-CRL-1739), Hs 746T (ATCC-HTB-135), KATO III (ATCC-HTB-103), NCI-N87 (ATCC-CRL-5822), SNU1 (ATCC-CRL-5971) and SNU5 (ATCC-CRL-5973) were obtained from the American Type Culture Centre. The human fetal gastric epithelial cell line GES-1 was provided by Professor Caiyun Fu (Zhejiang Sci-Tech University, China). All cells were tested for mycoplasma contamination prior to experimentation and intermittently tested thereafter by PCR. BJ-TERT cells were cultured in Dulbecco’s Modified Eagle’s Medium (DMEM) plus 20 mM HEPES, 17% Medium 199 (Gibco™ #11150067), 15% fetal bovine serum (FBS), and 1% GlutaMAX™ L-alanyl-L-glutamine dipeptide (Gibco™ #35050061). IMR90 cells were cultured in Eagle’s Minimum Essential Medium (EMEM) supplemented with 10% FBS, 5 mM sodium pyruvate (Gibco™, #11360070), 1% non-essential amino acids (Gibco™, #11140050), and 1% GlutaMAX™. GES-1, AGS, Hs746T and KATOIII were cultured in DMEM+20mM HEPES, 10% FBS and 1% GlutaMAX™. NCI-N87 and SNU-1 were cultured in RPMI +20mM HEPES, 10% FBS and 1% GlutaMAX™. SNU-5 was cultured in IMDM, 20%FBS and 1% GlutaMAX™.

### Plasmids, virus production and transduction

pBABE-puro was a gift from Morgenstern, Land and Weinberg (Addgene plasmid #1764) (Morgenstern & Land, 1990), pBABE-puro-myr-AKT1 and pBABE-puro-HRASG12V were described previously (Astle et al., 2012). HA-myrAKT1 was directly subcloned into pCW57.1 (Addgene plasmid #41393) to generate doxycycline-inducible pCW57.1-myrAKT1. The REBIR construct, a modified doxycycline-inducible mirE small hairpin RNA (shRNA) expression version of TRMPVIR, was a gift from Sang-Kyu Kim (Peter MacCallum Cancer Centre, Australia) by substituting Venus with enhanced blue fluorescent protein (EBFP2). The DharmaconTM GIPZ lentiviral shRNAs targeting human CBS gene were obtained from Horizon Discovery, UK and were tested for functional knockdown efficacy in BJ-TERT cells, and the two best shRNA hairpins were subcloned into the REBIR construct. 97-mer shRNA sequences are listed in Supplementary Table S3.

The cloning vector pBSK(+) containing the cDNA encoding human CBS isoform 1 was synthesised and purchase from Biomatik company (Biomatik Cooperation, Cambridge, Canada). CBS cDNA was subcloned into REBIR plasmid in which the dsRed2/mirE cassette has been removed and EBFP2 was replaced with the puromycin resistance gene. The resulting plasmid was designated pRT3-puro-CBS. HEK293T cells were used for virus production and viral transductions were performed as previously described.

### siRNA transfection of BJ-TERT and IMR-90 fibroblasts

The siRNA targeting CBS (#L-008617-00) and On-Target Plus Non-Targeting siRNA control (#D-001810-01) were purchased from Horizon Discovery, UK. siRNA transfection was performed by reverse transfection where cells were seeded onto plates containing transfection reagents and siRNA mixture. In 96-well plates, 0.1 μL (For BJ-TERT) or 0.4 μL (For IMR-90) Dharmafect 1 (Dharmacon-T-2001-01) and 20nM ON-TARGETplus SMARTPool siRNA per well were mixed and dispensed into each well. 4,000 BJ-TERT or 12,000 IMR-90 cells in a total volume of 100 μL per well was then added. The negative controls including cell suspension only (no Dharmafect, no siRNA) or Dharmafect only control (no siRNA) were also prepared. The fresh complete medium was replaced at 24 hours post-transfection.

### Protein analysis

Protein was extracted with SDS-lysis buffer (0.5 mM EDTA, 20 mM HEPES, 2% (w/v) SDS pH 7.9) and protein concentrations were determined with the Bio-Rad DC protein assay. Proteins were resolved by SDS-PAGE, transferred to PVDF membranes and immunoblotted with primary and horseradish peroxidase–conjugated secondary antibodies (Supplementary Table 2). The signals were visualized by Western-Lightning Plus ECL (Perkin-Elmer-NEL104001EA) and ChemiDocTM Imaging system (Biorad-17001401).

### Colony formation assay

Cells were seeded in 6-well culture plates. Media was refreshed every 2 days. At the end of experiments, cells were fixed with 100% (v/v) methanol for 30 mins at RT and then stained with 0.1% w/v crystal violet for 30 mins at RT. After intensively washing with H_2_O and drying plates, images were obtained using ChemiDocTM Imaging system. Total colony area, expressed as percentage of cell coverage per well, was determined using the ImageJ plugin Colony Area.

### Anchorage-independent soft agar assay

The anchorage-independent soft agar assay was performed as described by Borowicz and colleagues (Borowicz et al., 2014). Cells were seeded at density of 1×10^4^ per well in 0.4% (w/v) noble agar (Difco-214220) in 6-well plate with the basal layer of 0.6% w/v noble agar. 1 ml of culture medium in the presence or absence of doxycycline was added over the upper layer of agar and replaced twice weekly. Colonies were stained by 0.001% crystal blue for 30 minutes and then extensively washed with PBS. The number of sizable colonies (diameter > 50 um) were manually counted under transmitted light microscopy for quantitative analysis.

### Senescence-associated β-galactosidase staining

10μM EdU was added into the culture medium 24 hours prior to cell fixation with 2% (v/v) paraformaldehyde, 0.2% (v/v) glutaraldehyde for 5min at RT. Cells were then incubated with X-Gal staining solution (25% v/v citrate buffer, 5mM potassium ferrocyanide (Sigma-P3289), 5mM potassium ferricyanide (Sigma-P-8131), 150mM NaCl (MPBio, #0219484801), 2mM MgCl2 (Merk, #536092), 5mg X-Gal (Sigma, #B4252)) at 37°C for 24 hours. After permeabilisation with 0.5% (v/v) TritonX-100 (Sigma-T8532) in PBS, EdU was fluorescently labelled with Click-iT™ EdU AlexaFluor-488 imaging kit (Invitrogen, #C10337) according to the manufacturer’s instructions. Cells were counterstained with DAPI. Images were obtained using a fluorescence microscope Olympus BX-61 using a 20x objective. A minimum of 200 cells per sample were counted and the percentage of EdU or SAβGal positive cells are quantified. SAβGal was only counted as positive in the absence of EdU incorporation.

### H_2_S production measurement

H_2_S production was measured using 7-Azido-4-Methylcoumarin (AzMC) fluorescent dye (Sigma, #802409). The protocol was adapted from Szabo Laboratory with minor modifications (Modis et al., 2014). AzMC reaction master mix consisted of 200mM Tris-HCl pH8, 20mM L-Cysteine (Sigma, #30089), 1mM L-Homocysteine (Sigma, #69453), 100μM Pyridoxal 5’-phosphate hydrate (Sigma, #P9255), 20 μM AzMC in H_2_O was prepared and kept on ice. Cells were washed with cool PBS and then harvested in cell lysis buffer (50mM Tris-HCl pH8, 150mM NaCl, 1% v/v IGEPAL^®^ CA-630(SigmaAldrich-I3021), 1% v/v Triton-X100). Samples were kept on ice for 1 hour and then centrifuged at 20,000g for 10mins at 4°C before protein quantification by DC protein assay. 400 ug proteins were mixed with 100μl AzMC reaction master mix and incubated at 37°C in dark for 2 hours. Samples were read at 340nm excitation and 445nm emission wavelength using CytationTM3 Cell imaging multi-mode reader (BioTek).

### Gas chromatography-mass spectrometry (GC/MS) Metabolomics

After saline wash, cells were quenched by pouring liquid nitrogen into 6-well plates and then harvested with ice cold methanol:chloroform:scyllo-inositol (MeOH:CHCl3 9:1 v/v) containing 3μM scyllo-inositol as internal standard. The extracts were vortexed for 10 seconds and incubated on ice for 15 minutes. By centrifugation at 4°C for 3mins at 16,100g, the supernatant was collected and snap-frozen in liquid nitrogen and stored at −80°C.

The samples were evaporated to dryness by vacuum centrifugation. Prior to GC/MS analysis, samples were derivatised with 25μl 3% (w/v) methoxyamine in pyridine (Sigma, #226904/270970) for 60 min at 37°C with mixing at 750 rpm, followed by trimethylsilylation with 25μl BSTFA + 1 % TMCS (Thermo, #38831) for 60 min at 37°C with mixing at 750 rpm. The derivatized sample (1μl) was analyzed using Shimadzu GC/MS-TQ8040 system, running the Shimadzu Smart Metabolites NRM database, comprising approximately 475 metabolite targets. Statistical analyses were performed using Student’s t test following log transformation and median normalization. Metabolites were considered to be significant if their adjusted p-values after Benjamini-Hochberg correction were less than 0.05. Further data analysis and enrichment analysis were performed through MetaboAnalyst 4.0.

### Liquid chromatography-mass spectrometry (LC/MS) metabolomics

Thiol derivatization using N-Ethylmaleimide (NEM) and LC/MS based detection were described previously and optimized (Ortmayr et al., 2015). Cells were seeded four days before harvest. Cells were washed with ice cold Saline and then collected with ice cold extraction buffer containing 50 mM NEM (Sigma-Aldrich-E3876) in 80% v/v methanol and 20% v/v 10mM ammonium formate (Sigma-Aldrich-516961) at pH 7. Final concentration of 2 μM L-Valine-^13^C5,^15^N (Sigma-Aldrich-600148), D-Sorbitol-^13^C6 (Sigma-Aldrich-605514) and L-Leucine-^13^C6 (Sigma-Aldrich-605239) were added in the extraction buffer as internal standards. Samples were mixed at 4°C on a vortex mixer for at 1 hour at 1,000 rpm before centrifugation at 20,000 g for 10 mins at 4°C. Supernatants were stored on ice and processed as per described previously (Ortmayr et al., 2015). Briefly, metabolites were separated on an Dionex Ultimate 3000RS HPLC system (Thermo Scientific, Waltham, Massachusetts, USA) using an InfinityLab Poroshell 120 HILIC-Z (100 x 2.1 mm, 1.9 μm) HPLC column (Agilent Technologies, Santa Clara, California, USA) maintained at 25°C with buffer A (20 mM (NH_4_)_2_CO_3_, pH 9.0; Sigma-Aldrich) and solvent B (100% acetonitrile) at a flow rate of 300 μl/minute. Chromatographic gradient started at 90% B, decreased gradually over 10 min to 65% B, then to 20% B at 11.5 min, stayed at 20% B until 13 min, returned to 90% B at 14 min and equilibrated at 90% B until 20 min. Metabolites were detected by mass spectrometry on an Q-Exactive Orbitrap mass spectrometer (Thermo Scientific) using a heated electrospray ionisation source (HESI). LC-MS data was collected using full scan acquisition in positive and negative ion mode utilizing polarity switching. Data processing was performed using Tracefinder application for quantitative analysis (Thermo Scientific). NEM derivatised thiols were detected in negative ion mode. Metabolite identification was based on accurate mass, retention time and authentic reference standards. Peak intensities were normalised against internal standards and cell numbers, and the statistical analyses were carried out using one-way ANOVA.

### RNA-sequencing and analysis

RNA sequencing and analysis were described previously (Chan et al., 2020). Poly-A selective RNA-seq libraries were prepared using the TruSeq RNA sample preparation kit (Illumina^®^) and sequenced on Illumina^®^ NextSeq 500. HISAT2 (version 2.0.4) for the 75 bp single end reads, was used for alignment to the genome (hg19/GRCh37). Reads were counted using feature counts (version1.6.2) in Galaxy. The differential expression of genes was calculated utilizing the DESeq2 package v1.24.0 and plotted in R. Absolute gene expression was defined determining RPKM. Gene set enrichment analysis (GSEA) was performed according to the Hallmark gene set from the molecular signatures database MSigDBv6.1 (Broad Institute).

### Statistical analyses

Data is presented as mean ± SEM values for triplicate biological replicates as indicated in the figure legends. Data was presented using GraphPad Prism (8.1) and P-values < 0.05 were considered significant. Statistical analyses were not used to determine sample size.

## Acknowledgments

This work was supported by the National Health and Medical Research Council (NHMRC) of Australia (Project grants #1053792 and #1162052) and Cancer Council Victoria Project Grant #1184873. R.B.P was supported by Senior Research NHMRC Fellowship #1058586. E.S. is supported by a Victorian Cancer Agency Mid-Career Research Fellowship (MCRF19007). Z.H was supported by a Melbourne International Research Scholarship. We acknowledge the Centre for Advanced Histology and Microscopy at the Peter MacCallum Cancer Centre for their support of this work.

## Author contributions

J.K and R.B.P conceived the project with contributions from K.T.C and E.S. H.Z, J.K, K.T.C, X.H, S.B performed the experiments with contributions from X.W. and C.F. A.S.T. performed RNA-seq data analysis. D.A. and D.C performed LC-MS and D.P.D performed GC-MS. M.J provided the assistance for microscopy analysis. J.K, R.B.P, K.T.C and H.Z wrote the paper.

