## Supporting information for "Cystathionine-β-synthase is essential for AKT-induced senescence and suppresses the development of gastric cancers with PI3K/AKT activation"

### Supplementary Methods

#### Plasmid transfection, virus concentration & transduction

HEK293T cells were seeded 24 hours prior to transfection in tissue culture flasks at 80%-90% confluency. To generate the retrovirus, the transfection reagent master mixes were prepared by combining equal mass of plasmid DNA vectors, pEQ-PAM3-E, and RD114 envelope plasmid as previously describe (Gavrilescu and Van Etten, 2007). To generate the lentivirus, plasmid DNA vectors were combined with pMDL, pRSV-REV and pCMV-VSV-G packaging plasmids at the ratio of mass 3:1:1:1. The plasmid mixtures were combined with polyethylenimine (PEI, 5ug per ug plasmid), a cationic polymer for gene delivery. After vortex briefly, the Mixtures were incubated at RT for 25min and then added to the media in a dropwise manner. The culture media were refreshed at 24h post-transfection. At 48 and 54h post-transfection, the culture media containing viral particles was collected and passed through a 0.4μm filter. Virus-containing media were either concentrated and applied to target cells or stored at -80°C for later use.

To concentrate the media containing virus, ultracentrifuge tubes were filled with 25mL virus-containing media to top. The tubes were weighed and balanced (all need to be within 0.1 g) prior to ultracentrifuge spin in 70Ti rotor (Beckman-337922) at 25000g for 2h at 4°C (BECKMAN-OPTIMA L-100XP). Supernatants were discarded into 1% (w/v) Virkon™ S solution for disinfection. Ultracentrifugation was repeated twice and tubes were inverted to drain excess supernatant on paper soaked in 1% w/v Virkon™ S for 15 min. Pellets were carefully resuspended in 400μL DMEM. Concentrated virus was stored at -80°C until further use.

Cells were seeded 24 hours before virus infection at 60-70% confluency. The concentrated virus was added with 4μg/ml polybrene. 8 hours later, the second virus hit was performed. At 48 hours post-transduction, the media was removed and replaced with the complete media. The transduced cells were selected either by the defined antibiotics (e.g. 1μg/ml puromycin) or by cell sorting for the fluorescent marker.

#### CRISPR/Cas-9 gene deletion of PTEN

The PTEN sgRNA was designed using software provided at crispr.mit.edu. Cells were infected at 0.5 MOI with Cas9-expressing lentivirus and then sorted for mCherry fluorescence marker. Cells were then subsequently re-infected with lentiviruses expressing sgRNA and sorted for mCherry and GFP positivity.

#### AGS human gastric cancer xenograft

All animal experiments were performed with approval from the Animal Experimentation Ethics Committee at the Peter MacCallum Cancer Centre (Ethics number E626). Female NSG mice aged between 8-10 weeks were purchased from Peter MacCallum Cancer Centre Animal Facility, Australia. $5x10^6$AGS human gastric cancer cells transduced with RT3-CBS-puro in

100μl PBS-Matrigel mixture were implanted into the right flank of mice, using pre-cooled 0.3mL insulin syringes (BD #230-4533). When tumours reached an average volume of $100mm^3$ calculated by $V = (W^2 \times L)/2$, mice were randomized into two groups with one group administered with doxycycline both in the drinking water (0.2% w/v Doxycycline hyclate (Sigma # D9891) in 2% sucrose (Sigma #S8501)) and food (600 mg/Kg Doxycycline, SpecialtyFeeds #SF08-026). Mice were sacrificed once tumours reached $1200mm^3$.

Immunofluorescence staining

Cells were seeded at least 48 hours before fixation. 1 uM mitoTracker (Invitrogen #M22426) was added into the medium and incubated for 30 minutes at 37°C before fixation in 4% paraformaldehyde for 10 min at RT. After washing with ice-cold PBS, cells were incubated with the blocking buffer (5% v/v goat serum, 0.1% v/v Triton X-100 in PBS) for 30 minutes at 37°C. 100 ul the CBS antibody (Proteintech #14787-1-AP) diluted in the antibody dilution buffer (1% w/v BSA in PBS) at 1:100 was added onto the cells and covered with a coverslip. Cells were incubated at 4°C overnight. After washing with PBS at room temperature for 5 minutes of three times, cells were incubated with the secondary antibody EnVision+System-HRP labelled polymer goat-anti-rabbit (Dako-K4003) at room temperature for 30 minutes. Cells were washed in PBS at RT for 5 minutes of three times. The Fluorophore Opal520: Excitation 494 nm; Emission 525 nm, AKOYA #SKU FP1487001KT) diluted in the antibody dilution buffer at 1:100 was added onto the cells. After incubation at room temperature for 10 minutes, cells were washed in PBS at RT for 5 min of five times in dark. Cells were counterstained with DAPI 0.5 ug/ml in PBS for 5 minutes at room temperature followed by PBS wash prior to mounting with VECTASHIELD Antifade Mounting Medium. The images were visualized using Nikon C2 Confocal microscope with Nikon Plan Apo VC 60x oil immersion objective (NA 1.4, Nikon, Japan).

Tissue Microarray (TMA)

The construction of Gastric cancer patient TMA has been described previously (Wang et al., 2013). TMA slide composes of 120 tumour sections and 63 normal sections from 62 individual gastric cancer patients. These patients underwent gastrectomy from 2000 to 2005 in Changhai hospital, second Military Medical University, Shanghai, China. All patients have not received any anticancer therapy before surgery. The tissue samples were obtained with patient informed consent and the protocol was approved by Institutional Review Board of Second Military Medical University.

TMA slide embedded in paraffin was baked at 60°C for 1 hour before dewaxing. The slide was incubated in 100% v/v xylene for 3 mins of three times followed by 100% v/v ethanol for 1 min of three times, 70% v/v ethanol for 1 min and water wash for 1 min to complete dewax/rehydration procedures. Antigen retrieval was conducted in high pH buffer (Agilent-K8004) using pressure cooker for 45 mins. The slide was then washed in 0.1% v/v Tween 20 in TBS buffer for 5 mins and blocked in the blocking buffer (2% w/v BSA, 0.2% v/v TX-100, 1% v/v goat serum in PBS) for 1 hour at room temperature. The CBS antibody (Proteintech #14787-1-AP) diluted in the blocking buffer was applied on top of the slide and covered with coverslip. The slide was incubated at 4°C overnight before wash in 0.1% v/v TBST at RT for 5 min of three times. The secondary antibody EnVision+System- HRP labelled polymer goat-anti-rabbit (Dako-K4003) was applied followed by incubation at RT for 30mins. The slide was washed in 0.1% v/v TBST at RT for 5 min of three times. The Fluorophore Opal620 (Excitation 588 nm; Emission 616 nm; Cap Colour Amber) from Opal 7-Color Manual IHC kit (PerkinElmer-NEL811001KT) was diluted in 1X Plus Amplification Diluent (PerkinElmer-FP1498) and applied to the samples. The slide was incubated for 10 mins and washed in 0.1%

v/v TBST at RT for 5 min of five times in dark. It was then counterstained with DAPI and washed in PBS prior to mounting with VECTASHIELD Antifade Mounting Medium. The images were taken using VS120 Virtual Slide Microscopy (OLYMPUS-VS120-L100-W) and analysed using HALO™ Image Analysis Software (v2.2.1870.17, Indica Labs, Albuquerque, NM, USA). The same threshold settings were applied to individual patient sections. Total number of cells expressing cytoplasm CBS (Red fluorescence) was divided by total cell number (DAPI) from each section and was shown as %Red Positive. Average cytoplasm fluorescence intensity on tumour tissue section were compared to normal tissue section from the same patient and expressed as fold change relative to normal control.

Glutathione (GSH) assay

GSH assay was performed using Glutathione Assay Kit (Cayman Chemical #703002) according to the manufacturer's protocol. Briefly, cells were seeded 24 hours before harvesting in 10cm plate at 80% Confluency with equal cell number. One extra plate was prepared for cell number counting after cell harvesting. Cells were washed twice with PBS and lifted using rubber scraper in 1mL ice-cold PBS. After centrifugation at 400g for 5mins at 4°C, the supernatants were discarded and cell pellets were resuspended in 100µl MES buffer (50mM MES, 1mM EDTA, pH6). Samples were snapped frozen in liquid nitrogen and then thawed in the ice. The procedures were repeated twice for cell lysis. After centrifugation at 12,000 g for 15 min at 4°C, the supernatants (100 µl) were collected without disturbing the pellets. 100µl of 2.5M metaphosphoric (MPA, Sigma #239275) was added to the supernatants for deproteination. The solutions were then vortexed and kept at RT for 5mins prior to centrifugation at 3,000g for 2mins. 10µl of 4M triethanolamine (TEAM, Sigma #T58300) was added into the supernatants collected. The deproteinated and neutralized supernatants were used for the determination of the amount of total GSH by mixing with the assay reagents (provided from the kit) in a 96-well plate. After 25 min incubation in dark, the absorbance at 405 nm was measured using a Benchmark Microplate Reader. Data were normalised with cell number and GSH standard curve.

Bioenergetics analysis using the Seahorse XF96 Extracellular Flux Analyzer

All bioenergetics analyses were performed using the Seahorse Bioscience XF96 extracellular flux analyser (Seahorse Bioscience, Billerica, USA). Cells were seeded in the Seahorse XF96 96-well plate coated with Cell-Tek (3.5 ug/cm2, Corning). After incubation for the indicated time period, cells were washed with the assay media (unbuffered DMEM, 11 mM glucose, 2 mM glutamine, 1 mM sodium pyruvate, adjusted pH to 7.4 with 0.1 M NaOH) before incubation in 180 µl of the assay media and equilibrated in a 37°C non-$CO_2$ incubator for 1 hour. The assay protocol consisted of 3 repeated cycles of 3 minutes mixing and 3 minutes of measurement periods, with oxygen consumption rate (OCR) and extracellular acidification rate (ECAR) determined simultaneously. Basal energetics were established after three of these initial cycles, followed by exposure to the ATP synthase inhibitor, oligomycin (1 µM) for three cycles, then p-trifluoromethoxy- phenylhydrazone (FCCP, 1 µM), which uncouples oxygen consumption from ATP production, was added for a further three cycles. Finally, the mitochondrial complex III inhibitor antimycin A (0.5 µM) and the complex I inhibitor rotenone (0.5 µM) was added for three cycles. At the completion of each assay, the cells were stained with 10 µM Hoechst. Images were analysed using a Cellomics Cellinsight 1 to determine the cell number per well.

ROS detection

MitoSOX$^{TM}$ Red mitochondrial superoxide indicator (5 μM; Invitrogen #M36008) or 2′,7′-Dichlorofluorescein diacetate (DCFH-DA; 10uM; Sigma #35845) was added to cell culture and incubated at 37°C for 1 hour. Cells were trypsinized and harvested before analysis by Canto II.

Gene expression analysis

RNA isolation and purification were performed using the ISOLATE-II kit (Bioline #52073) according to the manufacturer's protocol. 500ng of purified RNA was treated with DNase (Promega #M6101) at 37°C for 15min followed by heat inactivation at 70°C for 15min. Complimentary-DNA (cDNA) synthesis by reverse transcription was performed using SuperScript$^{TM}$ III First-Strand Synthesis System per manufacturer's instruction under the following conditions: initial incubation at 37°C for 5min, reverse transcription by SuperScript$^{TM}$ III reverse transcriptase (Invitrogen #18080051), hexameric random primers and dNTPs at 47°C for 2 hours and deactivation at 70°C for 15min. Quantitative real time-PCR (qRT-PCR) reactions were performed using the StepOne Plus Real-Time PCR system (Applied Biosystems #4376600) with a +0.7°C melt-curve increment. Reactions were performed in triplicate using MicroAmp Optical 96-well plates (Applied Biosystems #N8010560) containing 8 ul cDNA sample, 10 ul v/v Fast SYBR green Master Mix (Applied Biosystems #4385612) and 0.1μM forward and reverse primers in 2ul. Changes in target gene expression were normalized to the non-POU domain-containing octamer-binding protein (NONO) housekeeping gene. Fold changes in gene expression were determined by $2^{(-\Delta\Delta Ct)}$.

Methylation specific PCR (MSP)

Cells were seeded 48 hours prior to genomic DNA (gDNA) extraction. Genomic DNA was extracted using NucleoSpin® Tissue Kit (Macherey-Nagel #740952) according to manufacturer's protocol. Cells were resuspended in lysis buffer with presence of 1.35mg/mL Proteinase K (Roche-RPROTKSOL-RO). Cell lysate was applied to DNA-binding column and gDNA was eluted after binding and wash silica membranes. DNA bisulphite modification was performed using EZ DNA methylation kit (Zymo Research #D5001) according to manufacturer's protocol. 500ng gDNA was incubated with CT conversion reagent for 16 hours at 50°C in dark, to allow non-methlylated Cytosine (C) residue to be converted to Uracil (U). gDNA was applied to DNA-binding column and desulphonated for 15mins at RT. Columns were then washed and gDNA was eluted in 10 μl M-Elution buffer. PCR condition and primer sequence were adapted from Zhao and colleagues (Zhao et al., 2012). PCR amplification reaction mixture was prepared in 100 μl aliquots containing 2 μl of bisulphite converted gDNA, 200 μM dNTPs (Roche #DNTPM-RO), 1 mM Primer, 1.5 mM $MgCl_2$, 50mM KCl, 10 mM Tris-HCl pH8.3 and 1.25 units GoTaq® DNA Polymerase (Promega #M3001). PCR amplification reaction was performed in the T100 thermal-cycler under the following conditions: initial denaturation at 95°C for 10min, followed by 35 cycles (94°C for 30s, 55°C for 30s and 72°C for 30s) and reaction deactivation at 70°C for 15min.

**Supplementary Figures**

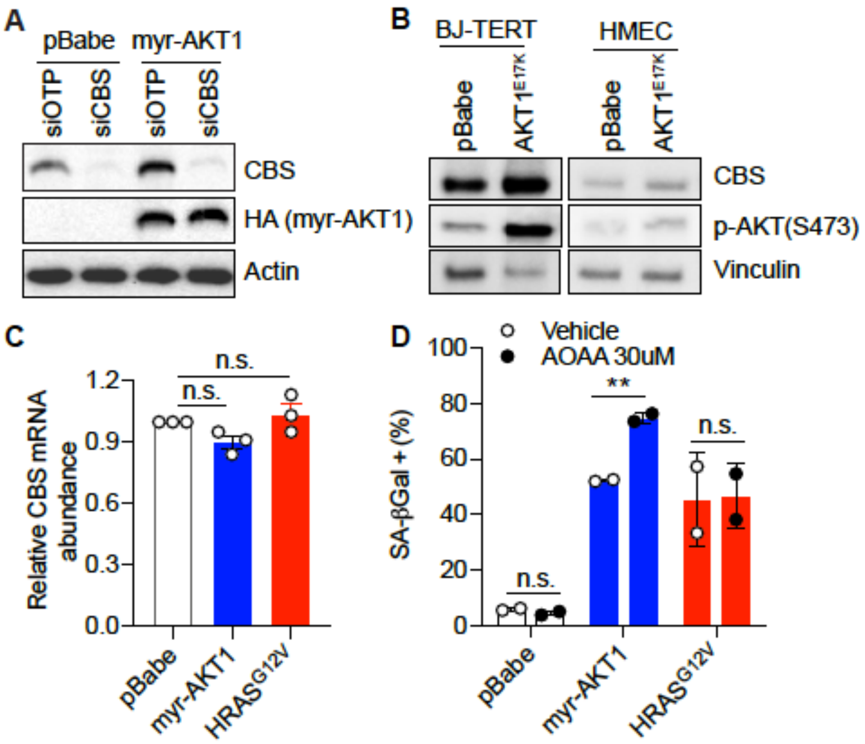


**Figure S1.** Increase of CBS expression and transsulfuration pathway activity in AKT-induced senescence. (**A**) IMR-90 human foetal lung fibroblasts were transduced with pBabe or myrAKT1. On day 5 post-transduction, cells were transfected with either CBS siRNA (siCBS) or control siRNA (siOTP). Western blot analysis of CBS and HA-tagged myrAKT1 on day 6 post-siRNA transfection. Actin was probed as a loading control. Representative of n=3 experiments. (**B**) BJ-TERT human skin fibroblasts and human mammary epithelial cells (HMEC) were transduced with either pBabe or $AKT1^{E17K}$. Western blot analysis of CBS and p-AKT at S473 expression on day 14 post-transduction. Vinculin was probed as a loading control. (**C**) BJ-TERT cells were transduced with pBabe, myrAKT1 and $HRAS^{G12V}$. The *CBS* mRNA expression level was measured on day 14 post-transduction by qPCR. Data are expressed as mean ± SEM (n = 3). n.s., not significant by one-way ANOVA. (**D**) BJ-TERT cells were transduced with pBabe, myrAKT1 and $HRAS^{G12V}$. Cells were treated with AOAA 30 μM or vehicle (DMSO) on day 5 post-transduction and quantification of the percentage of cells with positive staining for SA-βGal activity was performed after AOAA treatment for 6 days. Data are expressed as mean ± SD (n = 2). **, $P < 0.01$; n.s., not significant by student's t test.

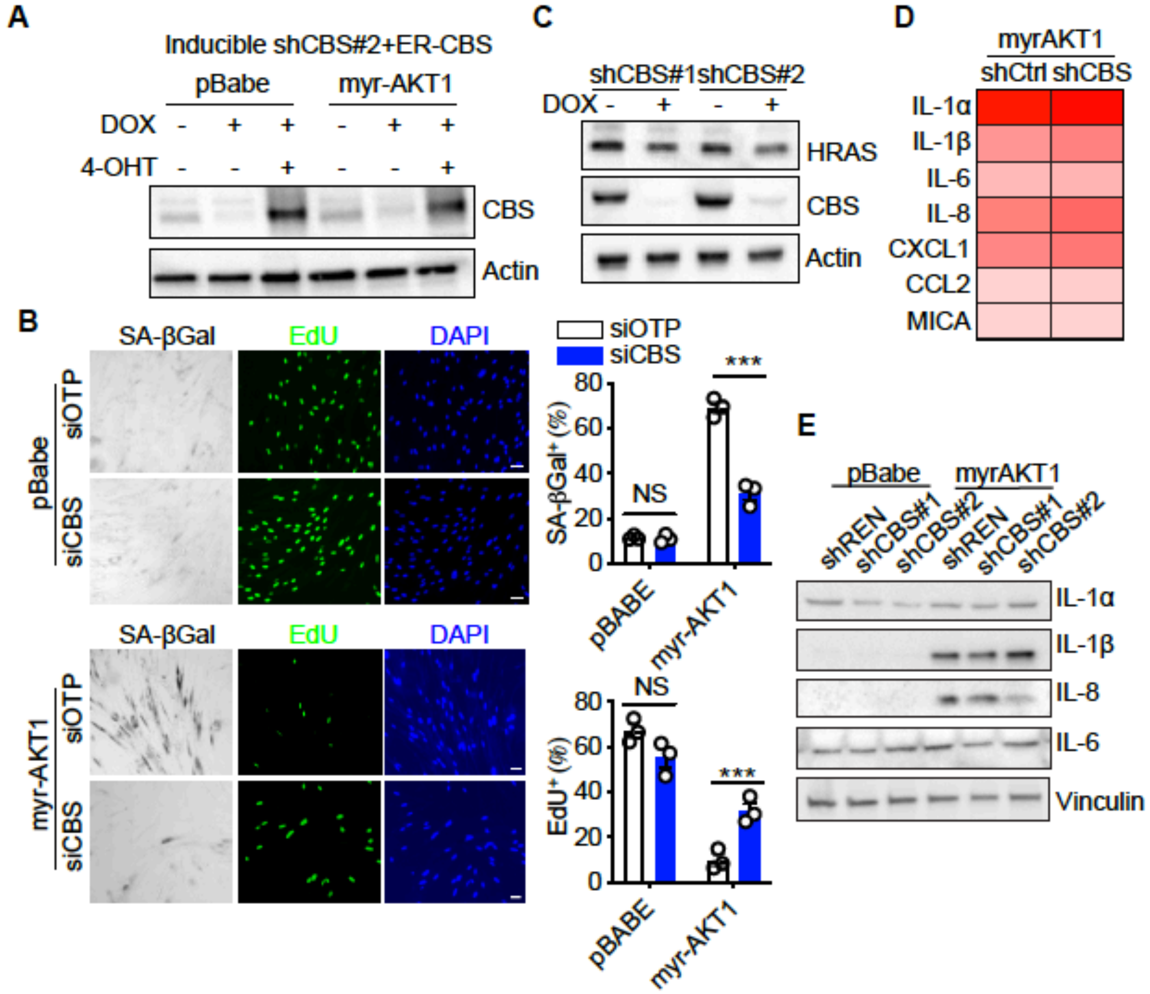


**Figure S2** Depletion of CBS promotes escape from AKT-induced senescence. (**A**) BJ-TERT cells expressing doxycycline-inducible CBS shRNA#2 and 4-OHT-inducible CBS were transduced with pBabe or myrAKT1, treated with doxycycline (1 μg/ml) ± 4-OHT (20 nM) on day 5 post-transduction. Western blot analysis of CBS expression on day 14 post-transduction. Actin was probed as a loading control. (**B**) IMR-90 human foetal lung fibroblasts were transduced with pBabe or myrAKT1. On day 5 post-transduction, cells were transfected with either CBS siRNA (siCBS) or control siRNA (siOTP). Images and quantification of percentage of cells with positive staining for SA-βGal activity and EdU were performed on day 6 post-siRNA transfection. Data were expressed as mean ± SEM (n = 3). ***, $P<0.001$; n.s., not significant by student's t test. (**C**) BJ-TERT cells expressing doxycycline-inducible CBS shRNA were transduced with $HRAS^{G12V}$, treated with doxycycline (1 μg/ml) on day 5 post-transduction. Western blot analysis on day 14 post-transduction. Actin was probed as a loading control. (**D**) BJ-TERT cells expressing myrAKT1 were transduced with pGIPZ-shCBS (myrAKT1-shCBS) or the control vector pGIPZ-NTC (myrAKT1-shCtrl) and the cells expressing pBabe transduced with the control vector pGIPZ-NTC (pBabe-shCtrl). The heatmap of mRNA expression levels of key SASP factors measured on day 14 post-transduction by qPCR. The data were normalised with *Actin* control and expressed as fold changes over pBabe-shCtrl cells. (**E**) BJ-TERT cells expressing doxycycline-inducible shCBS or control shREN were transduced with pBabe or myrAKT1, and then treated with doxycycline (1μg/ml) on day 5 post-transduction. Western blot analysis was performed on day 14 post-transduction. Vinculin was probed as a loading control. Representative of n=2 experiments.

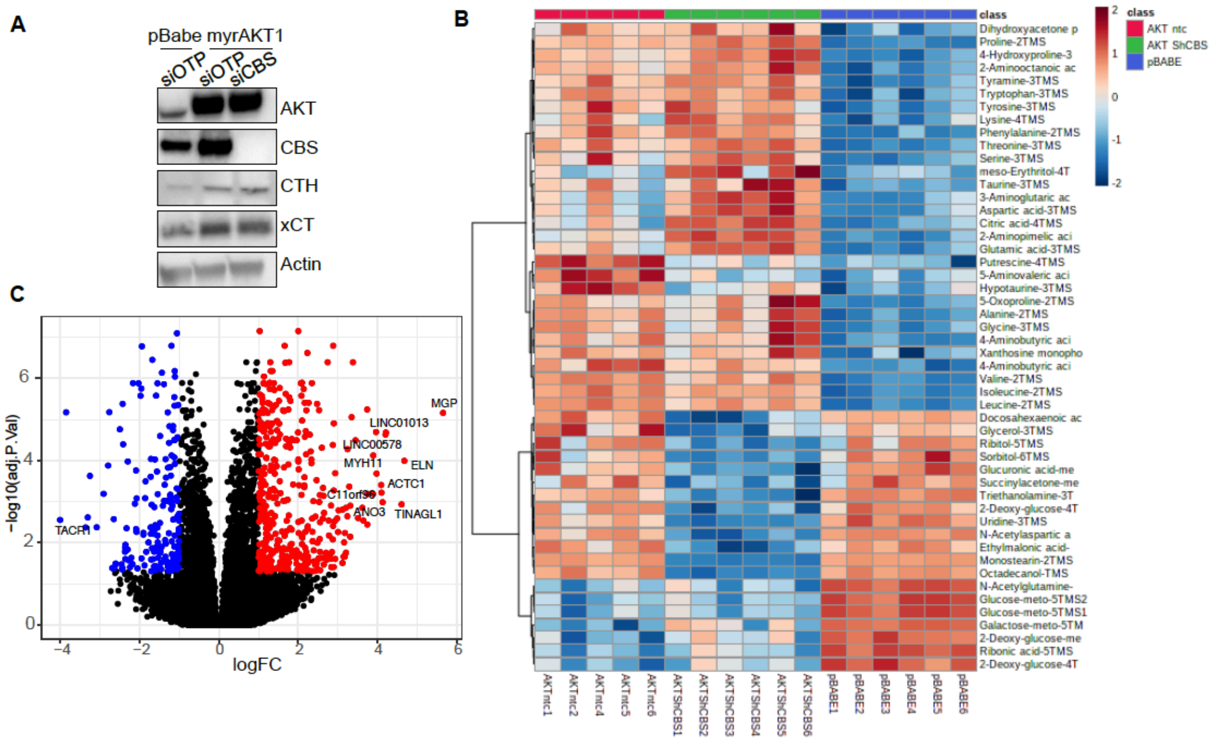


**Figure S3** Depletion of CBS in AIS cells does not alter activities of transmethylation and the transsulfuration pathways. (**A**) BJ-TERT cells were transduced with pBabe or myrAKT1. After four days cells were transfected with either CBS siRNA (siCBS) or control siRNA (siOTP). Western blot analysis of indicated proteins on day 6 post-siRNA transfection. Actin was probed as a loading control. (**B-C**) BJ-TERT cells were transduced with myrAKT1 followed by transduction with pGIPZ-shCBS (myrAKT1-shCBS) or control vector pGIPZ-NTC (myrAKT1-shCtrl) on day 6 post-transduction of myrAKT1. (**B**) The metabolic profiling by GC/MS was performed on day 14 post-transduction. n = 6 in both pBabe-shCtrl and myrAKT1-shCBS groups and n = 5 in myrAKT1-shCtrl group, with one sample excluded due to a technical issue in sample processing. The heatmap of top 50 metabolites analyzed by MetaboAnalyst 4.0. (**C**) RNAseq was performed on day 14 post-transduction. Volcano plot showing differentially expressed genes (FC ≤ -2 or FC ≥ 2, FDR ≤ 0.05) between cells expressing myrAKT1-shCBS vs. myrAKT1-shCtrl cells, which are downregulated (blue), upregulated (red) or not significant (black). The top 10 significantly upregulated and downregulated genes are indicated. n=3 biological replicates. (C)

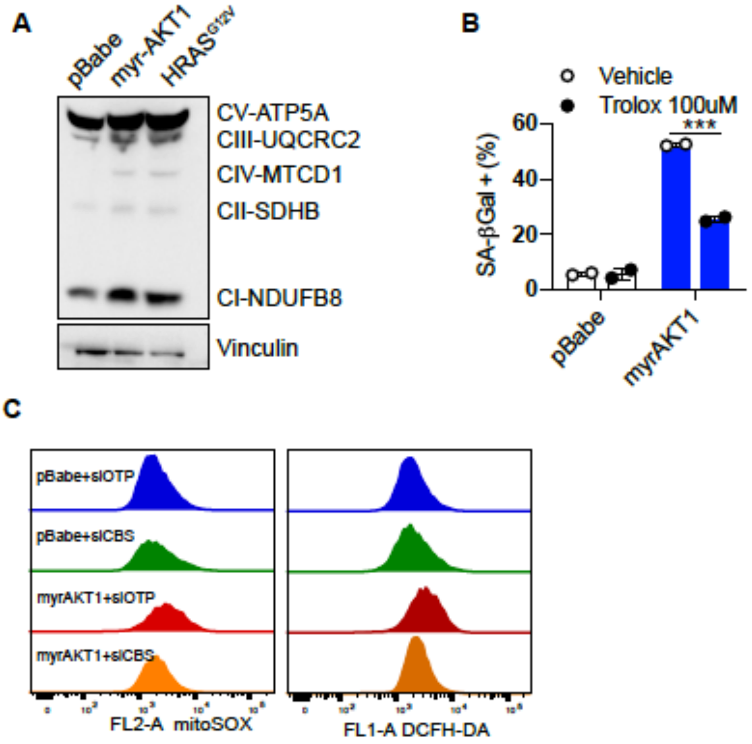


**Figure S4** CBS deficiency alleviates oxidative stress in AKT-induced senescent cells. (**A**) BJ-TERT cells were transduced with pBabe, myrAKT1 or $HRAS^{G12V}$. Western blot analysis of key complexes in the electron transport chain was performed on day 10 post-transduction. Vinculin was probed as a loading control. (**B**) On day 5 post-transduction, cells were treated with Trolox 100 µM or vehicle. Quantification of percentage of cells with positive staining for SA-βGal activity were performed after Trolox treatment for 6 days. Data were expressed as mean ± SD (n = 2). (**C**) BJ-TERT cells were transduced with either pBabe or myrAKT1. After five days cells were then transfected with either CBS siRNA (siCBS) or control siRNA (siOTP). Flow cytometric analysis of the mitochondrial superoxide production by MitoSOX and the cytoplasmic ROS production by $H_2$DCFDH-DA on day 6 post-siRNA transfection.

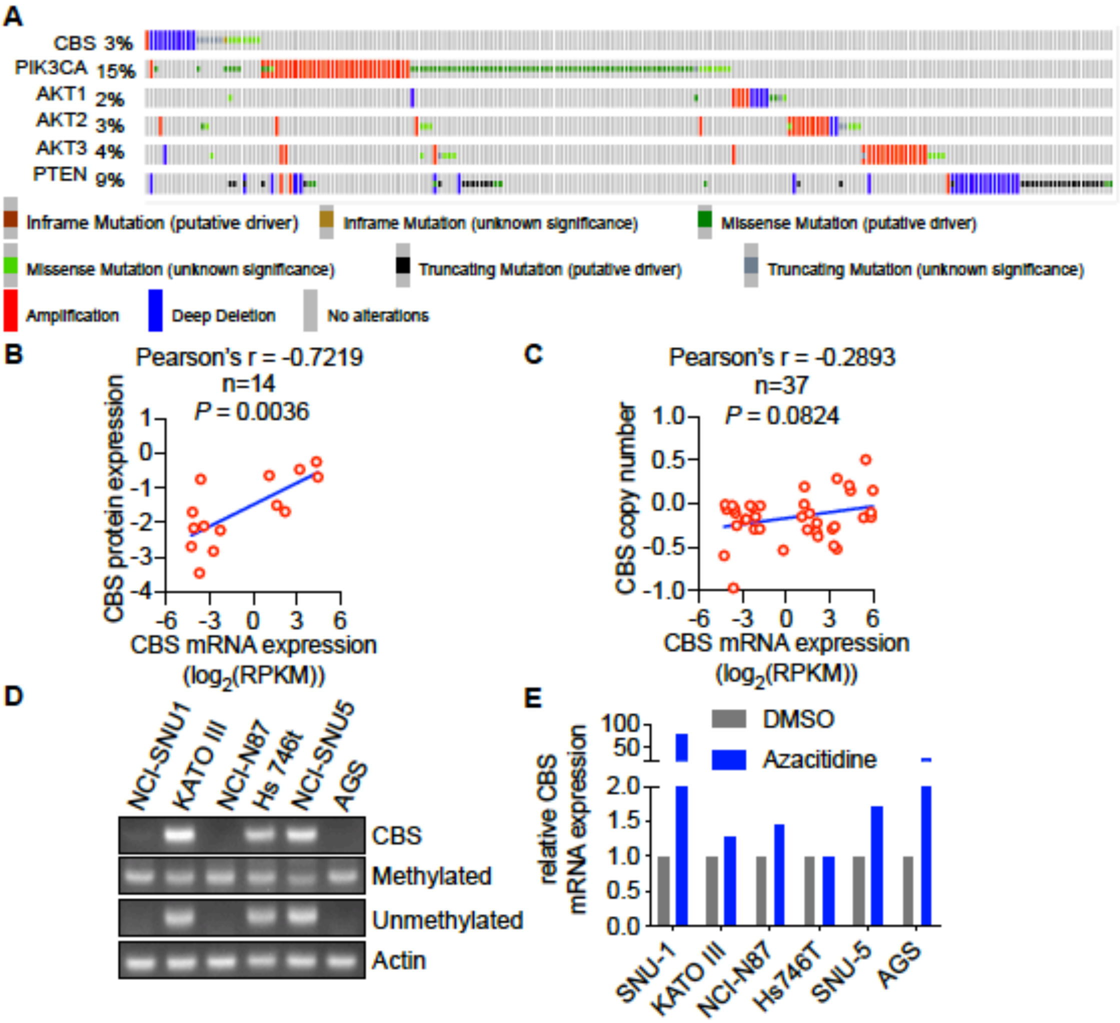


**Figure S5** CBS expression is suppressed in tumour tissues and human cell lines of gastric cancer. (**A**) CBS, PIK3CA, AKT1, AKT2, AKT3 and PTEN gene expression in 777 gastric cancer patient samples from 5 studies (OncoSG, 2018; Pfizer and UHK, Nat Genet 2014; TCGA, Firehose Legacy; U Tokyo, Nat Genet 2014; UHK, Nat Genet 2011) using the cBioPortal (http://www.cbioportal.org) (**B**) Correlation of *CBS* mRNA expression with CBS protein expression in 14 gastric cancer cell lines and (**C**) Correlation of CBS mRNA expression with CBS copy number in 37 gastric cancer cell lines based on the data retrieved from the Cancer Cell Line Encyclopaedia. (**D**) The methylation in CBS promoter region of six gastric cancer cell lines was examined by methylation-specific PCR and visualized after agarose gel electrophoresis. Actin was used as a loading control. (**E**) Gastric cancer cells were treated with 2 µM Azacitidine for 48 hours. *CBS* mRNA expression levels were analysed by qRT-PCR with the fold changes relative to the untreated controls.

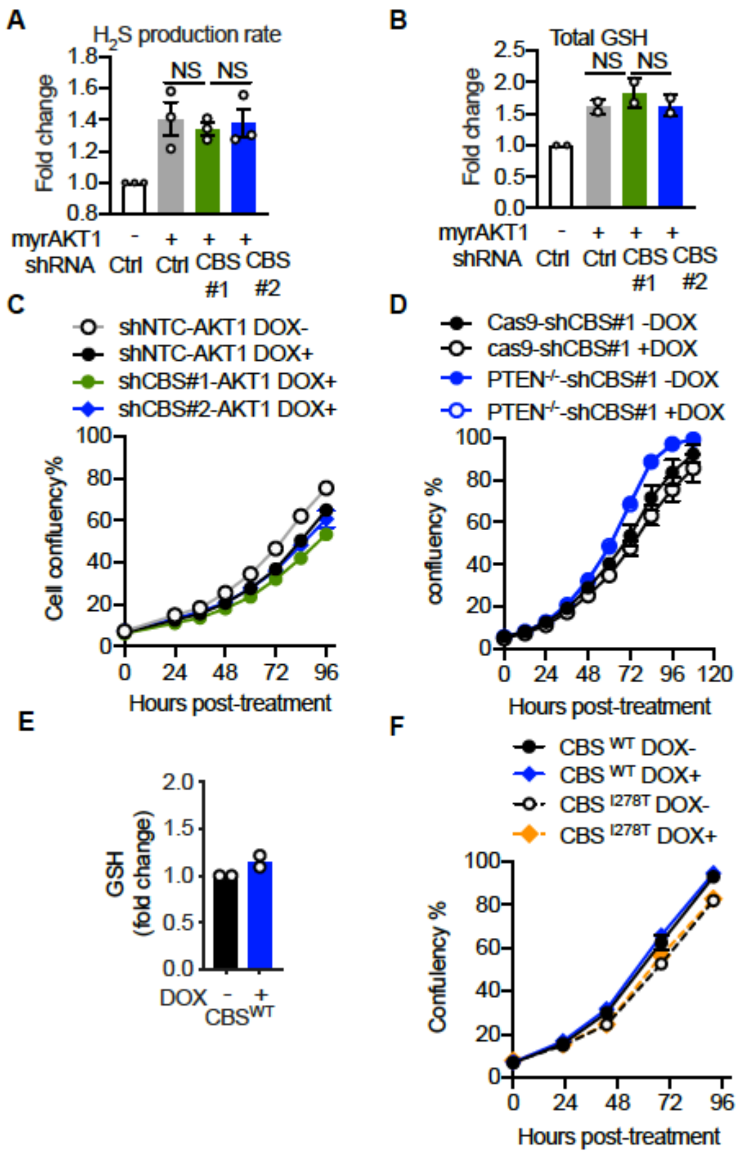


**Figure S6** Loss of CBS synergizes with PI3K/AKT pathway to promote gastric cancer pathogenesis. (**A-C**) GES-1 gastric epithelial cells were stably transfected with doxycycline-inducible myrAKT1 and pGIPZ-shCBS or control pGIPZ-NTC (shCtrl). Cells were treated with doxycycline (0.75 µg/ml). (A) $H_2S$ production. (**B**) total GSH level were measured on day 3 post-doxycycline induction. Data were expressed as mean ± SEM (n = 3) for (A) and mean ± SD (n = 2) for (B). (**C**) Cell confluency was measured by IncuCyte. Data were expressed as mean ± SEM (n=3). (**D**) GES-1 cells with PTEN knockout by CRISPR or Cas9 control were transduced with doxycycline-inducible CBS shRNA#1. Cells were treated with doxycycline (0.75 µg/ml). Cell confluency was measured by IncuCyte. Data were expressed as mean ± SEM (n=4). (**E-F**) AGS gastric cancer cells were stably transfected with doxycycline-inducible $CBS^{wt}$ or $CBS^{I278T}$ and treated with doxycycline (0.08 µg/ml for $CBS^{wt}$ or 1 µg/ml for $CBS^{I278T}$). (**E**) GSH abundance was measured on day 3 post-doxycycline induction and expressed as the relative fold changes. Data are expressed as mean ± SD (n = 2). (**F**) Cellular confluency was measured by IncuCyte. Data are expressed as mean ± SEM (n = 3).
